# Restoration decisions in highly variable systems: Whether, when, and how to enhance kelp

**DOI:** 10.64898/2026.10.28.762303

**Authors:** Mei Z. Blundell, Marissa L. Baskett

**Affiliations:** Department of Environmental Science and Policy, University of California Davis, Davis, California, USA

**Keywords:** Mathematical model, Simulations, Restoration, Decision-making, highly variable ecosystems, *Nereocystis leutkeana*, *Macrocystis pyrifera*

## Abstract

In highly variable ecosystems, natural variability can obscure signals of decline that warrant intervention, as there is often a lack of long-term data needed to distinguish between declines that are transient and likely to recover naturally and those that would be permanent without intervention. Implementing restoration for a species involves deciding on a threshold condition at which to intervene, duration that this threshold needs to be met before intervening, life stages at introduction, and amounts to introduce. Current bull and giant kelp restoration efforts in California were spurred by persistent kelp loss that started in 2013 following a disease outbreak among predators of the primary herbivore and a marine heatwave. We asked how decisions about intervention cue, threshold, wait time, life stage at introduction, and amount affect population outcomes against a backdrop of environmental variability. We analyzed additional context-dependencies on grazing and climate intensity. To address these questions, we built and parameterized a stochastic stage-structured model for bull and giant kelp, in which we simulated kelp enhancement approaches under a range of grazing and climate scenarios. In a global sensitivity analysis of kelp enhancement decisions, we found that grazing intensity and life stages at introduction were the most influential determinants of kelp biomass and persistence outcomes. The kelp enhancement approach involving temporarily introduced juveniles restored the germ banks of bull and giant kelp, but the two species differed in their responses to that intervention, highlighting that optimal intervention approach is species-dependent.

Waiting longer for natural recovery before intervening reduced total intervention effort but also reduced biomass recovery potential. These results support empirical findings from kelp enhancement pilot projects in California and highlight context-dependencies that inform restoration decision-making in kelp forests and other highly variable ecosystems.

## Introduction

### Restoration in highly variable systems

Accounting for a system’s natural variability is an acknowledged challenge in restoration, often because there is a lack of long-term data needed to distinguish between declines that are part of natural variability and those that warrant intervention (Holl and Aide, 2011; Scheffer et al., 2001; Suding and Hobbs, 2009). This affects management decisions for intervention cue, threshold, timing, effort, and approach.

Natural variability can mask relevant cues and thresholds which signal a need for intervention. To illustrate with an example from forest management, bark beetle populations irregularly but continuously cycle between periods of low population size and periods of outbreak during which their large populations lead to widespread tree mortality (Lorio, 1984; Singh et al., 2024). Knowledge gaps in the drivers of this irregular cycling make it challenging to identify cues for when the next outbreak will happen, predict how intense and long-lasting an outbreak will be, and ultimately to make informed decisions about a threshold condition at which to intervene (Biedermann et al., 2019; Marini et al., 2017).

Faced with the possibility of recovery without intervention in highly variable systems, managers can save resources by delaying intervention in case natural recovery occurs, i.e., “passive restoration” and “wait-and-see” approaches (reviewed in Gann et al., 2019). On the other hand, delaying intervention too long can have ecological consequences, such as organisms adapting to a “degraded” state and challenging an eventual shift to a target state (Chaparro Pedraza et al., 2021), erosion of recovery capacity over time due to cumulative stress (Osborne et al., 2017), and accumulation of an ecosystem’s “recovery debt” in organism abundance, species diversity, and nutrient cycling (Moreno-Mateos et al., 2017).

Beyond decisions on whether and when to intervene, natural variability also affects decisions regarding intensity of intervention effort and the types of intervention to use, especially as appropriate or optimal types of intervention might change over time. Due to biophysical conditions continuing to fluctuate after an initial action, managers might choose to invest in continuous or long-term approaches to achieve certain goals, e.g., maintaining a target abundance or species composition (Cayton et al., 2023). For bark beetle management, uncertainty in the drivers, duration, and intensity of outbreaks constrains decisions on how much effort to allocate to different interventions: removing infested trees to slow the outbreak, killing beetles with insecticides or natural enemies, using antiaggregation pheromones to prevent aggressive beetle behavior, or reducing tree density to improve forest-level resistance (reviewed in Fettig et al., 2022). Each approach acts in the system differently, so decisions about how to allocate limited resources among them offer strategic opportunities to influence intervention outcomes.

Navigating decisions on whether, when, and how to intervene in highly variable systems further depends on life history and disturbance regime. Species longevity affects the rate of population recovery after disturbance, with short-lived species recovering faster than longer-lived species (Capdevila et al., 2022; White et al., 2022).

Thus, managers might be more inclined to delay intervention and wait for natural recovery if the focal species is short-lived, or might choose to extend their wait time for longer-lived species if the costs of delayed recovery are acceptable. Additionally, critical life stages that form natural demographic buffers, e.g., dormant early life stages (hereafter referred to as “germ banks”), can especially influence population growth rate and thus be effective targets for intervention (Crouse et al., 1987; Doak et al., 1994; Morris et al., 2024).

Restoration decisions in naturally fluctuating systems are exacerbated by uncertainty in the perturbations associated with climate change. Shifts in climate regime affect recovery potential, and consequently might influence the intensity of intervention effort that is appropriate (e.g., grazing intensity in rangelands amid fluctuating drought, (Scasta et al., 2023)), the feasibility of waiting for natural recovery (e.g., unlikely natural recovery of palatable grasses over decades under drier climate, Cipriotti et al., 2019), and the prioritized intervention approaches (e.g., “climate-ready interventions” such as stress-priming and selective breeding, (Bernath-Plaisted et al., 2025) and adaptive management strategies (Derner et al., 2023)). In some cases, release from anthropogenic or biotic stressors can improve recovery potential, and thereby reduce intervention effort and necessity (Morris et al., 2024; Zengel et al., 2021). For example, grazer exclusion in highly variable desert grasslands led to significant recovery of vegetation, soil nutrients, and species diversity within ten years, and reduced reliance on active planting for restoration (Hou et al., 2024; Miao et al., 2015). Understanding the relative importance and context-dependencies of each decision can help managers and practitioners prioritize their data collection and decision analysis (Canessa et al., 2015).

### Kelp system and restoration

Kelp is an example system where natural variability and exacerbating uncertainties related to climate change affects all the above intervention decisions. Kelp forests are dynamic ecosystems that fluctuate temporally and spatially between forested and urchin barren states (Dayton et al., 1984; Steneck and Johnson, 2013; Fig 1). Recent persistent kelp declines are motivating restoration globally (Eger et al., 2022). Current considerations for kelp restoration include the following (summarized in unpublished manuscript led by Baskett and Pinsky): Is decline in adult biomass, visible from the surface, an effective cue for intervention? Or would earlier cues such as low juvenile recruitment or shift in urchin grazing behavior lead to more effective restoration (Ling and Keane, 2024; Reeves et al., 2022)? How long can intervention wait for natural recovery or clearer signals of persistent decline (personal communication via Science Advisory Committee to CDFW Kelp Restoration and Management Plan)? Is the scale of kelp enhancement (i.e., supplying kelp material to the environment) needed to reach a target population size feasible (McHugh et al., 2025; Reed et al., 2024)? Is the scale of grazer suppression needed to enable kelp recovery feasible (Arroyo-Esquivel et al., 2023; McHugh et al., 2025; public communication from Giant Giant Kelp Restoration Project)? What kelp enhancement strategies, i.e., what life stages at introduction, are most effective for restoration and how can their efficacy be improved (see Fig 2; Eger et al., 2022; Graham et al., 2023; McHugh et al., 2025)? Understanding the efficacy of intervention cues, thresholds, wait times, effort, and life stages at introduction will help clarify which decisions and their context-dependencies especially influence kelp restoration outcomes. Exploring these questions can help inform restoration decision-making in other naturally highly variable systems.

**Figure 1:**
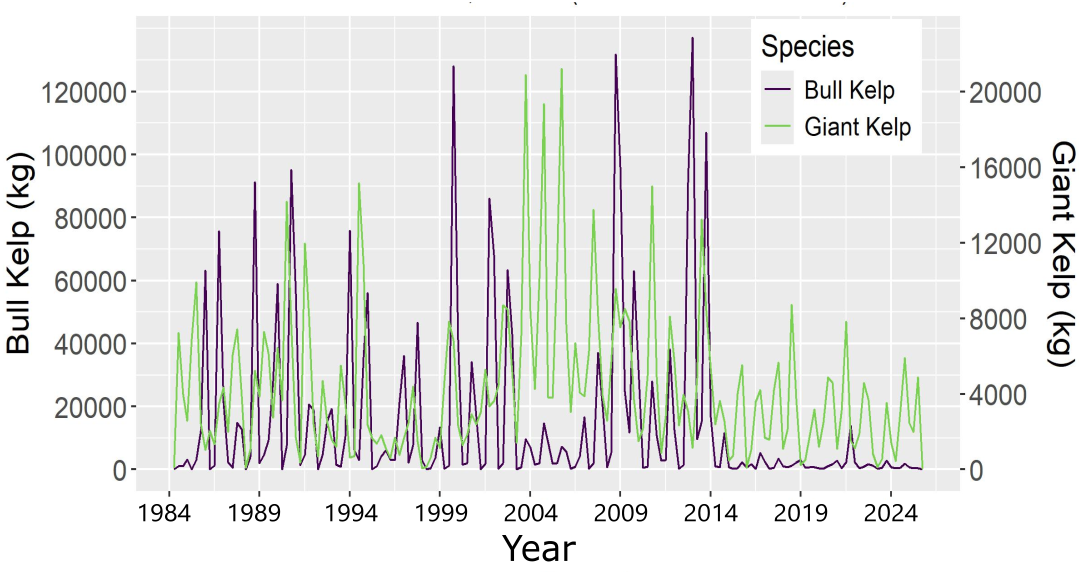
Empirical time series. Time series (1984-2025) of bull kelp (purple) and giant kelp (green) biomass estimated from Landsat 5,7, and 8 kelp cover data (Bell et al., 2024) and scaled up to a hypothetical 100*m* x 100*m* kelp bed. Bull kelp data are from 37.8 to 42 °N. Giant kelp data are from 32.53 to 35.45 °N. See *Landsat biomass estimates* subsection of *Appendix: Additional methods* and GitHub repository for estimation approach.

**Figure 2:**
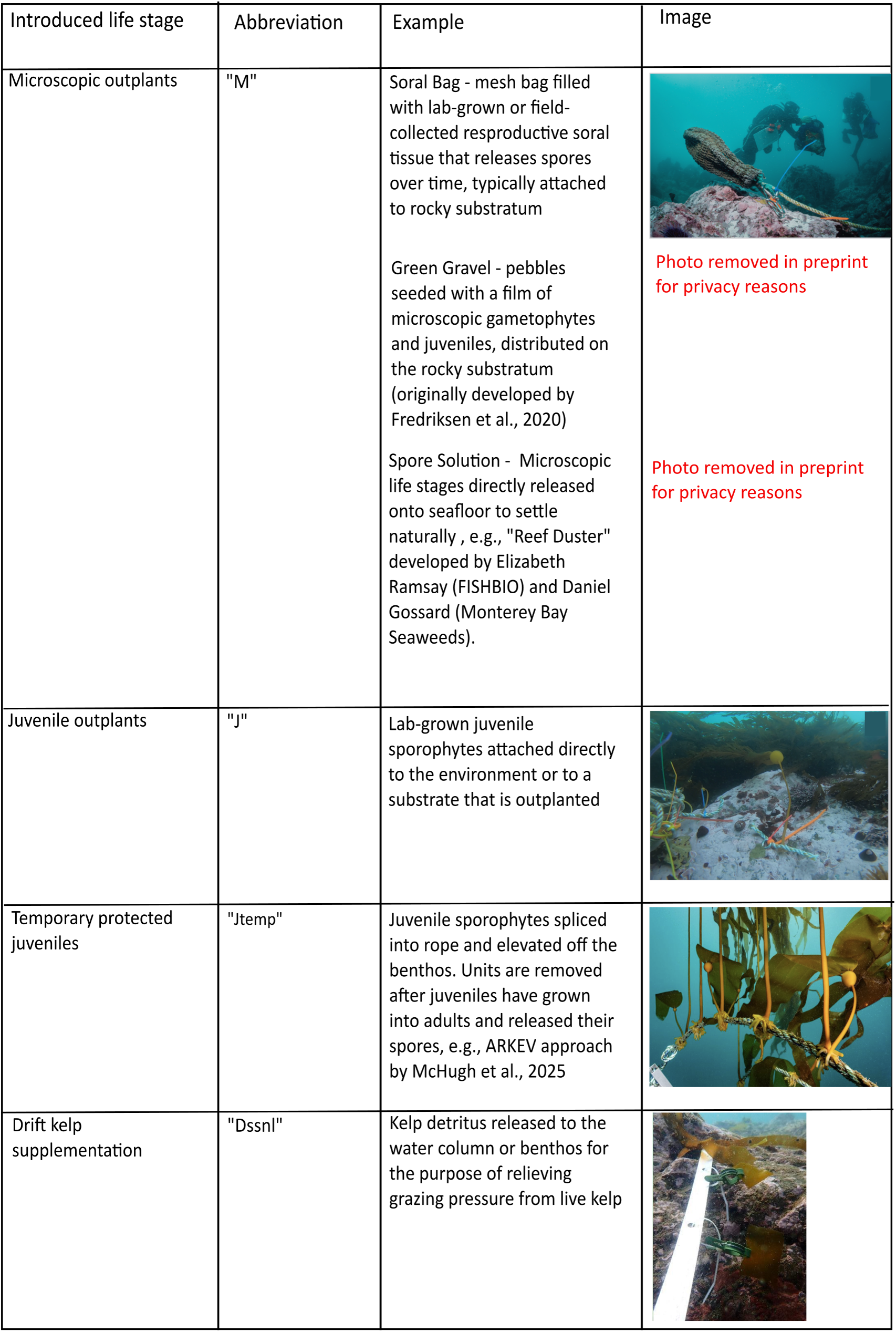
Kelp enhancement interventions. Summary of kelp life stages that are introduced to the environment for kelp enhancement. Photo credits from top to bottom: Patrick Webster (Fig 4A in Graham et al., 2023); Bennett Bugbee (Fig 4B in Graham et al., 2023); Patrick Webster; Bennett Bugbee and Tristin McHugh (Fig 30 in McHugh et al., 2025).

### Paper overview

Here we quantify how, in a naturally highly variable system, the decisions of intervention cue, threshold of decline, wait time, amount of introduced material, and life stages at introduction affect population outcomes, with kelp as our example. We compare across contexts to understand how population outcomes depend on the interactions between those decisions and biophysical attributes: species life history, grazing intensity, and climate scenario. In particular, we address the following questions: (1) Which intervention decisions and biophysical attributes are most influential for population persistence and biomass in a 10,000 *m*^2^ kelp bed? (2) Which life stages at introduction are most effective for restoring population persistence and biomass, and do they differ between annual and perennial species? (3) How does delaying intervention by waiting for natural recovery impact population persistence and biomass, and do those relationships depend on species, grazing intensity, and climate scenario? To address these questions, we construct a stochastic stage-structured model that tracks biomass through the life stages of the annual bull kelp (*Nereocystis leutkeana*) and perennial giant kelp (*Macrocystis pyrifera*), from which we simulate interventions spanning the decision space under a range of grazing rates and “good” and “bad” year frequencies, and summarize resulting biomass and persistence outcomes.

## Methods

### Study system

Kelp demography follows a seasonal pattern. For bull kelp and giant kelp, recruitment of microscopic gametophytes to the juvenile sporophyte stage can occur in all seasons, but typically peaks in early spring when upwelling of nutrient-rich water brings favorable conditions for recruitment. Density-dependent effects include an Allee effect, wherein gametophyte fertilization occurs within 1 *mm* (Boland et al., 1983; Reed, 1990), and negative density dependence due to competition for light and space with adults and other juveniles (Dayton et al., 1984, 1999; Dean et al., 1989; Reed, 1990).

Spring and summer for both species see growth from juvenile to adult sporophytes.

Adult sporophyte biomass is generally greatest at the end of summer, as most individuals reach their maximum height at this time of year (Reed et al., 2008; Schiel and Foster, 2015; Springer et al., 2010).

In the fall, adult sporophytes release spores from reproductive packets called sori. While fecundity is greatest in late fall and early winter for both species in California, giant kelp can reproduce year-round (Anderson and North, 1966; Buschmann et al., 2006; Neushul, 1963; Reed et al., 1996).

Kelp mortality is especially high in the winter due to storms that can rip away whole individuals (Foreman, 1970; Johnson and Koehl, 1994; Reed et al., 2011). Juvenile sporophytes are less susceptible to storm-driven mortality due to their short heights and structural properties (Dobkowski and Crofts, 2021; Dobkowski et al., 2019).

Throughout the year, but especially during winter storms, blade and frond tissue break off and exist in the system as drift kelp (Rassweiler et al., 2025). Drift kelp has a deterrent effect on the grazing of the other kelp life stages, as urchins, the dominant grazers for both species, prefer to consume drift over live kelp (Randell et al., *in press*; Duggins, 1983; Rennick et al., 2022; Sheridan and Randell, 2026). However, when urchins are starving, they will swarm together to actively attack kelp holdfasts, which leaves swaths of reef devoid of kelp, i.e., urchin barrens (Harrold and Reed, 1985).

Many kelp forest-urchin barren systems demonstrate hysteresis: kelp population recovery requires much lower grazing than that which originally led to population collapse (Ling et al., 2015).

In addition to the seasonal patterns described above, kelp demography also fluctuates with year-to-year environmental variability, especially with water temperature and nutrient content. Recruitment (Deysher and Dean, 1986; Hernández-Carmona et al., 2001), growth (Fales et al., 2023), and fecundity (Carney and Edwards, 2010; Weigel et al., 2023) increase when cool, nutrient-rich water is brought to the surface by upwelling. Warm water temperatures above 18° C increase gametophyte mortality (Weigel et al., 2023) and breakage due to temperature-driven weakening of tissues (Fales et al., 2023; Zimmerman and Kremer, 1984), while decreasing blade growth (Supratya et al., 2020) and recruitment (Deysher and Dean, 1986; Hernández-Carmona et al., 2001).

### Model overview

We used a stage-structured matrix model with a seasonal timestep to track transitions of biomass through microscopic germ bank (*M*), juvenile sporophyte (*J*), adult sporophyte (*A*), and drift kelp (*D*) life stages. We chose to track biomass rather than number of individuals for convenience of parameterizing with canopy cover data from Landsat, and to accommodate the roles of high biomass turnover and breakage at the adult stage. To capture germ banking and adult age structure, we incorporate several age classes within the microscopic and adult stages (Fig 3). The kelp demographic dynamics play out against a backdrop of stochastically occurring “good”, “neutral”, and “bad” years, which are inspired by strong upwelling, no significant upwelling or marine heatwaves, and marine heatwaves, respectively. We parameterized the model based on California bull kelp and giant kelp. Using model simulations, we (1) conducted a global sensitivity analysis (Harper et al., 2011) to identify the most influential intervention decisions and biophysical attributes for population persistence and biomass, (2) compared the efficacy of different life stages at introduction across species, and (3) showed how intervention wait time varies with population persistence and biomass outcomes for both species under a range of grazing intensities and climate scenarios.

**Figure 3:**
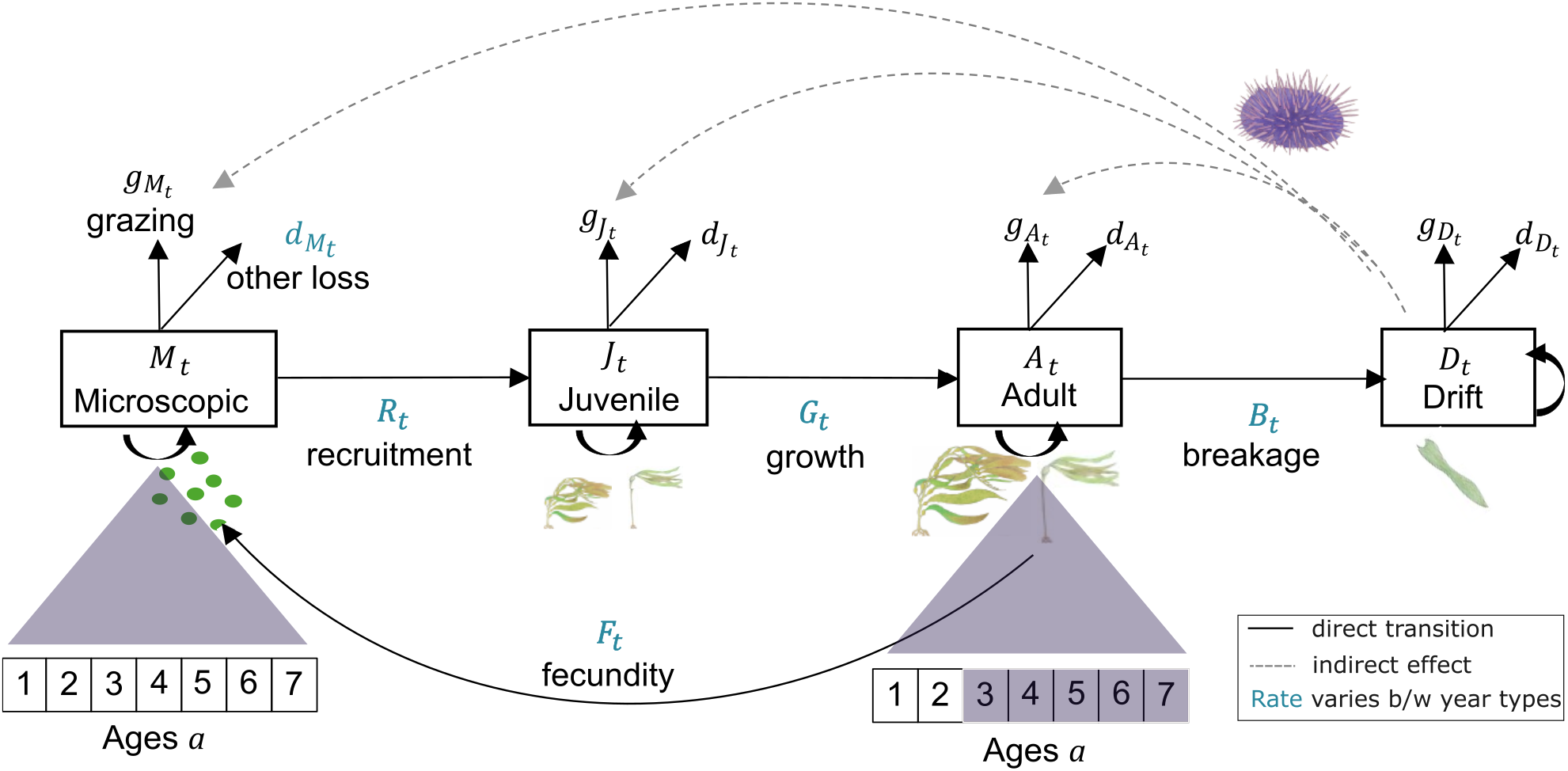
Box model diagram. Biomass moves through microscopic *M_t_*, juvenile *J_t_*, adult *A_t_*, and drift kelp *D_t_* stages from season to season. Transition rates with subscript *t* vary seasonally. Those that are in teal blue text also vary between ”good”, ”neutral”, and ”bad” years. Solid black arrows represent direct transfers of biomass. Dotted gray arrows represent the indirect effect of drift kelp biomass on biomass in other life stages due to grazer satiation. Age classes 3-7 of the adult stage are shaded to indicate the case of minimal adult age structure in bull kelp (small 2nd year cohort is possible for bull kelp, mortality rates allowing).

### Demographic dynamics

#### Microscopic stage survival

Early kelp life stages include spores, gametophytes, eggs, sperm, and microscopic sporophytes. We combine these stages into a single category *M* in our model, because the transitions which typically last days or weeks between them are much faster than the months-long transition to identifiable sporophytes *J* (Lü ning and Neushul, 1978; Reed et al., 1992; Schiel and Foster, 2006), although see Edwards (2022) for a review of delays in microscopic stage transitions and the possibility of kelp germ banks.

Before negative density dependence, the microscopic stage biomass of age *a* at time *t* which remains in *M*, 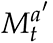*^′^*, is that which does not recruit at rate *r*, get grazed at rate *g_Mt_*, or otherwise die at rate *d_Mt_* :

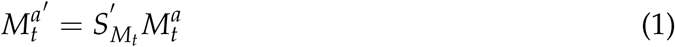

where

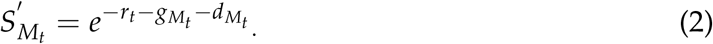

For survival and transition terms involving microscopic, juvenile, and adult life stages, we included negative density dependence in the saturating Beverton-Holt style to represent competition for light and space. Although it is dubious whether gametophytes are abundant enough in the field to generate negative density dependence (about 1-10 per *mm*^2^ from Reed et al., 1991), we found that omitting negative density dependence at all stages produced unrealistic chaotic dynamics in test runs. Therefore, we used small effect size *β_M_* to represent weak shading and crowding for the microscopic stage *M*. After negative density dependence from concurrent *M*′_*t*_ and newly produced microscopic stages *A″_t_* the amount of *M* biomass that remains by the next season is

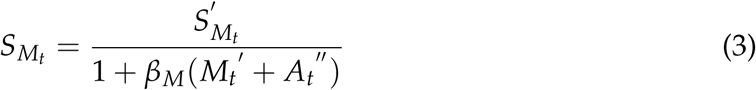

where total existing microscopic biomass before shading and crowding is

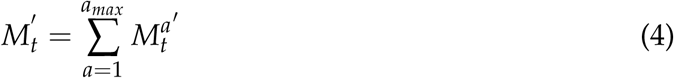

and total newly produced microscopic biomass before shading and crowding is

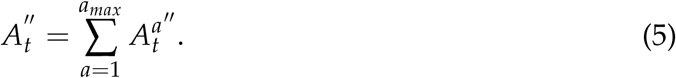

See Fecundity subsection below for calculation of newly produced microscopic biomass from age *a* adults, 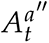.

#### Recruitment

Microscopic biomass enters the sporophyte recruitment process at rate *r_t_* and is subject to grazing and non-consumptive mortality at rates *g_Mt_* and *d_Mt_*, respectively. We include an Allee effect, at strength *α*, due to the recruitment requirement for high density (Boland et al., 1983; Reed, 1990). The microscopic recruits grow to juvenile size with growth factor *γ_Jt_* . Altogether, the amount of new juvenile sporophyte biomass before negative density dependence is

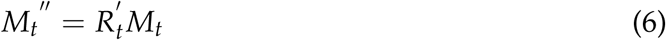

where

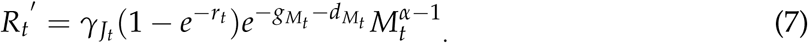

These new juveniles then face negative density dependence *β _JA_* due to competition with other juveniles *J_t_^′^* and adults *A_t_^′^* already present, along with fellow new juveniles *M_t_^′′^* and new adults *J_t_^′′^* from the current seasonal transition (reviewed in Schiel and Foster, 2006). As above, we also include the negative density dependence *β_M_* coming from microscopic biomass. Altogether, the proportion of biomass entering the juvenile stage is

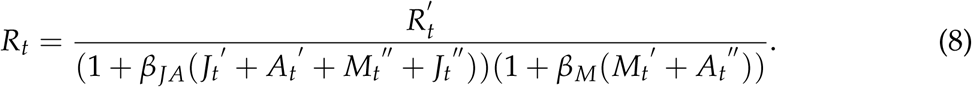

See Fig S1 for a plot of density-dependent recruitment as parameterized in this model.

#### Juvenile survival

Biomass stays in the juvenile stage and survives between time steps if it does not mature at rate *m_t_*, get grazed at rate *g_Jt_*, or otherwise die at rate *d_Jt_* . *S_Jt_*′, the proportion of remaining biomass in the juvenile stage before density-dependence, is

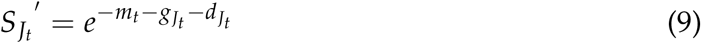

and the amount of remaining biomass is

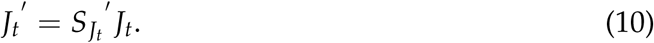

Juveniles also face negative density dependence *β _JA_* due to competition with other juveniles *J_t_*′ and adults *A_t_*′, as well as new recruits *M_t_*″ and new adults *J_t_*″ from the current seasonal transition. After this shading and crowding, remaining proportion of juveniles is

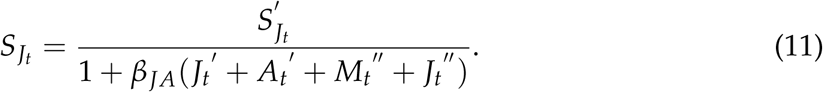

#### Growth

Juvenile biomass moves to the adult stage with maturation rate *m_t_*and experiences grazing at rate *g_Jt_* and non-consumptive mortality at rate *d_Jt_* . Maturing biomass grows by factor *γ_A_*. Before density-dependent effects, the amount of juvenile biomass that transitions to the adult stage is

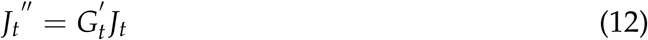

where

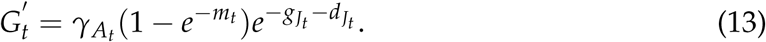

New adults face negative density dependence *β _JA_* due to competition with juveniles *J_t_*′ and adults *A_t_*′ already present, along with new recruits *M_t_*″ and fellow new adults *J_t_*″ from the current seasonal transition. After this shading and crowding, realized growth is

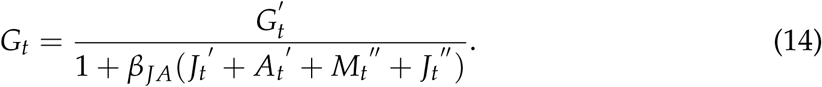

#### Adult Survival

In addition to loss due to grazing at rate *g_At_*, adult sporophytes of age *a* face non-consumptive mortality during time step *t* at rate 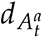. This mortality represents loss of whole individuals, e.g., due to wave action, rather than senescence-related breakage of living tissue at rate *b_t_* (see Breakage subsection below). Adult sporophytes grow back tissue at rate *w_t_*during the season. To represent fecundity, a small proportion *f_t_* of biomass leaves the adult stage as released spores. Taken together, the amount of remaining age *a* adult biomass before negative density dependence during time step *t* is

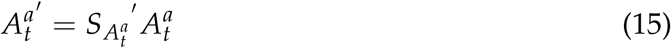

where

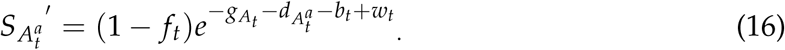

Summed across all age classes, the total biomass remaining in the adult stage between seasons before density dependent effects is

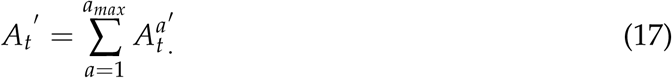

The remaining adults then experience negative density dependence *β _JA_* due to competition with juveniles *J_t_*′ and other adults *A_t_*′ already present, as well as new recruits *M_t_*″ and new adults *J_t_*″ from the current seasonal transition. The proportion of adult biomass remaining in each age class becomes

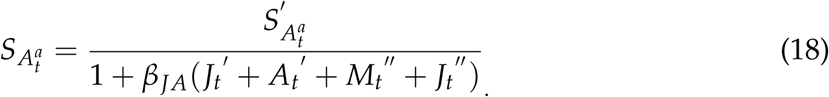

#### Fecundity

The proportion of adult biomass released as spores during time step *t* is *f_t_*. Of this amount, the proportion that survives grazing at rate *g_Mt_* and mortality at the first-year rate 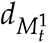 is

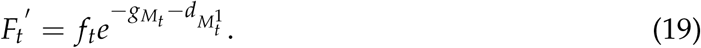

This means the total biomass of new microscopic stages before density dependence is

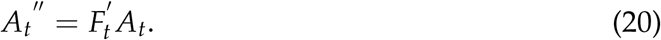

After competition of strength *β_M_* for light and space with already present microscopic stage biomass *M_t_*′ and concurrent new biomass *A_t_*″, the proportion of adult biomass that enters the first-year microscopic cohort 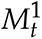 is

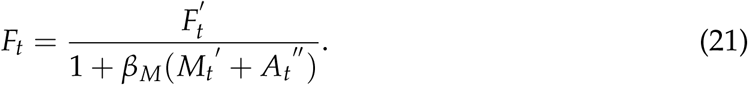

#### Breakage and drift kelp

We model breakage, the creation of drift kelp, from two sources: a proportion *ɛ* of kelp tissue released as “crumbs” from grazing, and tissue that has broken away at rate *b_t_* from living adult sporophytes due to wave action or tissue senescence. The proportion of age *a* adult biomass that becomes drift kelp during time step *t* and is not released as spores (1 *− f_t_*), grazed at rate *g_At_*, or otherwise lost at rate 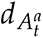 is

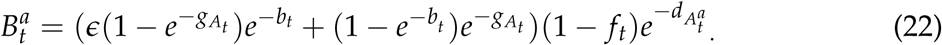

Drift kelp leaves the system due to grazing at rate *g_Dt_* and export at rate *d_Dt_*, such that the remaining proportion between seasonal time steps is

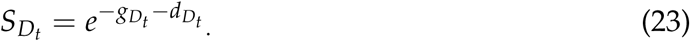

#### Grazing

For simplicity, we model grazing behavior without explicitly including the population dynamics of the dominant urchin herbivore. With this approach, we miss a feedback between kelp and grazer abundance, but we do represent different levels of grazing intensity (as proxy for abundance) with “low”, “medium”, and “high” grazing rates *g*. We justify our modeling choice under the assumption that grazer behavior rather than abundance drives kelp population dynamics (Harrold and Reed, 1985; Karatayev et al., 2021), although see Behrens and Lafferty (2004) for a counterexample.

To model the behavioral switch in urchin grazing, we define *E_at_* as a proportion of the fixed overall grazing rate *g* that is allocated to kelp life stage *a* at time *t*, such that *E_Mt_* + *E_Jt_* + *E_At_* + *E_Dt_* = 1. The proportion of grazing effort allocated to drift kelp, *E_Dt_*, initially increases at rate *k* with the available biomass of drift kelp, *D_t_*, and asymptotically saturates at *E_Dt_* = 1:

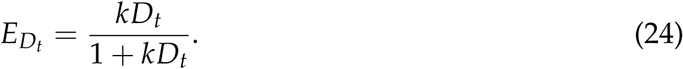

Grazers allocate their remaining effort 1 *− E_Dt_* to the other life stages according to their preferences for microscopic stages *p_M_*, juvenile sporophytes *p_J_*, and adult sporophytes *p_A_*, such that

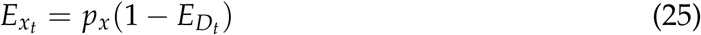

for *x ∈ M*, *J*, *A* and *p_M_* + *p_J_* + *p_A_* = 1. We assume that grazers’ preference for each stage is constant. The grazing rates on each life stage are then

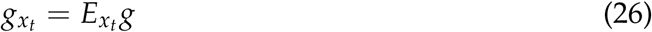

for *x ∈ M*, *J*, *A* (Fig S2).

#### Matrix model

We consolidate the above-described dynamics into a matrix of transitions between all stages and ages (details in *Matrix model* subsection of *Appendix: Additional methods*). In this process, we “age up” biomass in the germ bank and adult stage at the end of winter each year by shifting the survival terms 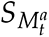 and 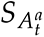 down to the subdiagonal of the transition matrix whenever *t ∈ winter*.

### Environmental dynamics

The demographic dynamics described above play out against a backdrop of stochastically occurring “good”, “neutral”, and “bad” years, inspired by strong upwelling (good) and marine heatwaves (bad). In “bad” years, microscopic mortality *d_Mt_* (Weigel et al., 2023) and breakage *b_t_* due to temperature-driven weakening of tissues are greater (Fales et al., 2023; Zimmerman and Kremer, 1984), while recruitment *r_t_* (Deysher and Dean, 1986; Hernández-Carmona et al., 2001) and growth factors *γ_Jt_* and *γ_At_* (Supratya et al., 2020) are lower relative to the “neutral” year parameterizations. We assume that in the long run, and fecundity *f_t_* is also reduced by “bad” years, though see Le et al. (2022). We make these effects in the opposite direction for “good” years.

To set the probabilities of “good”, “neutral”, and “bad” years, we created a set of scenarios based on possible climate futures. We used sea surface temperature (SST) projections from the NOAA-GFDL GFDL-ESM4 model prepared for CMIP6, as proxy for climate impacts on kelp, where nutrient availability and temperature are inversely correlated drivers of kelp population dynamics (Fales et al., 2023; Garćıa-Reyes et al., 2014; Leichter et al., 2023; Weigel et al., 2023; Zimmerman and Kremer, 1984). We selected projections at 122 °W and 36 °N in Monterey Bay, because this location is within the ranges of both bull and giant kelp. We calculated year-type probabilities for our “historical” scenario from the historical hindcast of the GFDL-ESM4 model (Krasting et al., 2018), and for our “mild”, “moderate”, and “extreme” scenarios from shared socio-economic pathways SSP126, SSP370, and SSP585 respectively (John et al., 2018a). We used a rolling climatology window approach to account for acclimatization and adaptation, such that “bad” and “good” years are defined by at least 3 months of anomalously high and low SST compared to the past 30 years (Logan et al., 2014). See details in *Environmental dynamics* subsection of *Appendix: Additional methods*.

### Kelp enhancement interventions

This paper explores five decisions for kelp enhancement: cue, threshold, wait time, effort, and introduced life stages. Intervention cue represents a choice among three signals of kelp decline: summer adult biomass, spring juvenile biomass, and grazing behavior, measured as the difference between proportion of drift and adult biomass consumed in a season (Table 1). This approach allows us to explore a few signals of decline that are potentially management-relevant cues (Ling and Keane, 2024; Reeves et al., 2022).

**Table 1:** Summary of intervention decision and biophysical context variables.

| **Variable** | **Type** | **Values** |
|---|---|---|
| species | biophysical context | bull kelp<br>giant kelp |
| climate scenario | biophysical context | historical<br>mild ∼ SSP126<br>moderate ∼ SSP370<br>extreme ∼ SSP585 |
| grazing intensity | biophysical context / intervention decision | low ∼ 10% grazed<br>medium ∼ 50% grazed<br>high ∼ 90% grazed |
| cue | intervention decision | adult biomass<br>juvenile biomass<br>grazing behavior |
| threshold | intervention decision | 0.0–0.1, percentile |
| wait time | intervention decision | 0–10, integers |
| effort | intervention decision | 0%–100% of maximum effort |
| life stages at introduction | intervention decision | NONE ∼ no intervention<br>M ∼ microscopic outplants<br>J ∼ juvenile outplants<br>Jtemp ∼ temporary protected juveniles<br>Dssnl ∼ drift supplement<br>MJ ∼ microscopic and juvenile outplants<br>MJtemp ∼ microscopic outplants and temporary protected juveniles<br>MDssnl ∼ microscopic outplants and drift supplement<br>JDssnl ∼ juvenile outplants and drift supplement<br>JtempDssnl ∼ temporary protected juveniles and drift supplement<br>MJDssnl ∼ microscopic and juvenile outplants, and drift supplement<br>MJtempDssnl ∼ microscopic outplants, temporary protected juveniles, and drift supplement |

Intervention threshold is a percentile of the distribution of cue values from one hundred simulations without intervention in the “historical” climate scenario. To illustrate, if we choose “summer adult biomass” as the cue and 0.1 as the percentile (as in Fig 4D), then intervention is “flagged” whenever the simulated summer adult biomass dips below the 10th percentile of summer adult biomass from the final years of 100 historical simulations. We “unflag” intervention whenever the threshold is no longer met, either due to successful intervention or natural recovery.

**Figure 4:**
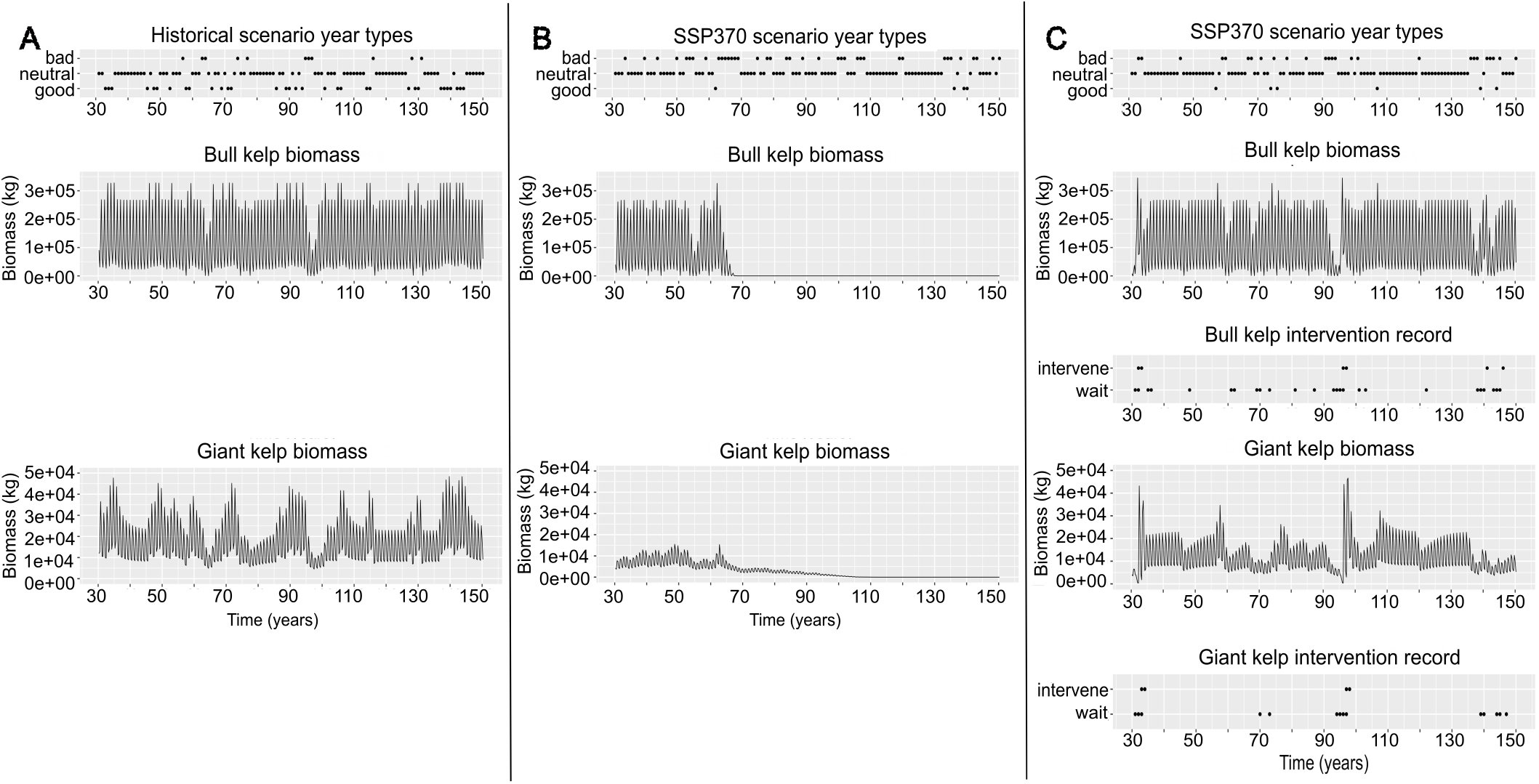
Simulated time series. Top rows in panels A, B, and C show stochastically generated time series of year types used for the corresponding simulations. A) Simulated time series of bull kelp and giant kelp biomass in the ”historical” climate and ”low” grazing scenario without intervention. B) Simulated time series of bull kelp and giant kelp biomass in the ”moderate” (SSP370) climate and ”low” grazing scenario without intervention. D) Simulated time series of bull kelp and giant kelp biomass in the ”moderate” (SSP370) climate and ”low” grazing scenario with annual intervention effort of 500kg temporary protected juveniles ”Jtemp”, 3 years of wait time, adult biomass cue, and percentile threshold of 0.1 for bull kelp and 0.05 for giant kelp (ARKEV-like intervention).

Wait time is the number of years that intervention must be “flagged” before it is implemented. We start counting years whenever intervention is “flagged” anew and stop counting whenever intervention is no longer needed, or when the wait time is met and intervention begins. This approach accommodates a naturally highly variable system where variation is common and managers might wait to intervene until there is a clear indicator of persistent decline or loss.

Effort is the percent of a maximum amount of microscopic or juvenile biomass to introduce to the system. To illustrate, if we choose an intervention approach that involves microscopic and juvenile outplanting (“MJ”, see Fig 2) at 90% effort, then 0.9 *∗ e f f M_max_* kg will be added to 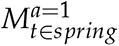 and 0.9 *∗ e f f J_max_* kg will be added to *J_t∈spring_* during intervention years.

We model the following life stages at introduction and their combinations: microscopic outplants “M”, juvenile outplants “J”, temporary protected juveniles “Jtemp”, and drift kelp supplements “Dssnl” (Table 1, Fig 2). Kelp outplanting typically occurs as an annual event in the spring (Eger et al., 2022), which we capture in the model by adding introduced “M”, “J”, and “Jtemp” biomass to the stage distribution vector at the beginning of spring in intervention years. Restoration practitioners can discourage urchins from grazing live kelp by engaging their fear response with predator introduction (e.g., sea stars, rock crabs, otters) and by satiating them with drift kelp supplementation (Randell et al., *in press*; Eger et al., 2022; Kriegisch et al., 2019; McHugh et al., 2025; Rennick et al., 2022; Sheridan and Randell, 2026). Here, we explore the possibility of drift supplementation as a frequent seasonal effort to prolong grazer deterrence.

### Parameter estimation

We chose parameter values through a combination of literature search and tuning, i.e, sweeping through ranges of possible values and selecting those that produced biologically reasonable simulations. We aimed to calibrate model output with adult biomass estimated from Landsat (Fig 1, Bell et al., 2026), as well as adult bull kelp abundance data (Stekoll et al., 2006), juvenile giant kelp abundance data (Nelson et al., 2021; Reed et al., 1997), and adult giant kelp abundance data (Rassweiler et al., 2025). Due to limitations of satellite imagery, Landsat data can only provide information about canopy cover and omits estimation of juvenile and understory kelp abundance (Reed et al., 2026). However, it covers a broader spatial range and is more accessible than *in situ* surveys, and thus provides a decent rough estimate for adult kelp biomass at a spatial scale relevant for this model. Informed by these sources, we targeted a 50,000 to 200,000 kg range for adult bull kelp biomass, and a 10,000 to 50,000 kg range for adult giant kelp biomass in our modeled 10,000*m*^2^ kelp beds. When data were not available from the literature to inform seasonal values of demographic parameters, we adjusted around literature-derived means to produce the seasonal dynamics described in the *Study system* subsection above: high juvenile biomass after spring recruitment, high adult biomass after summer and fall growth, minimum adult biomass after winter storms, and high microscopic biomass after fall and winter spore release. We parameterized intervention effort by scaling up existing kelp enhancement trials in California (McHugh et al., 2025). See Table 2 for a complete list of parameter values and sources. Further details are in *Parameter estimation* subsection of *Appendix: Additional methods*.

**Table 2:**
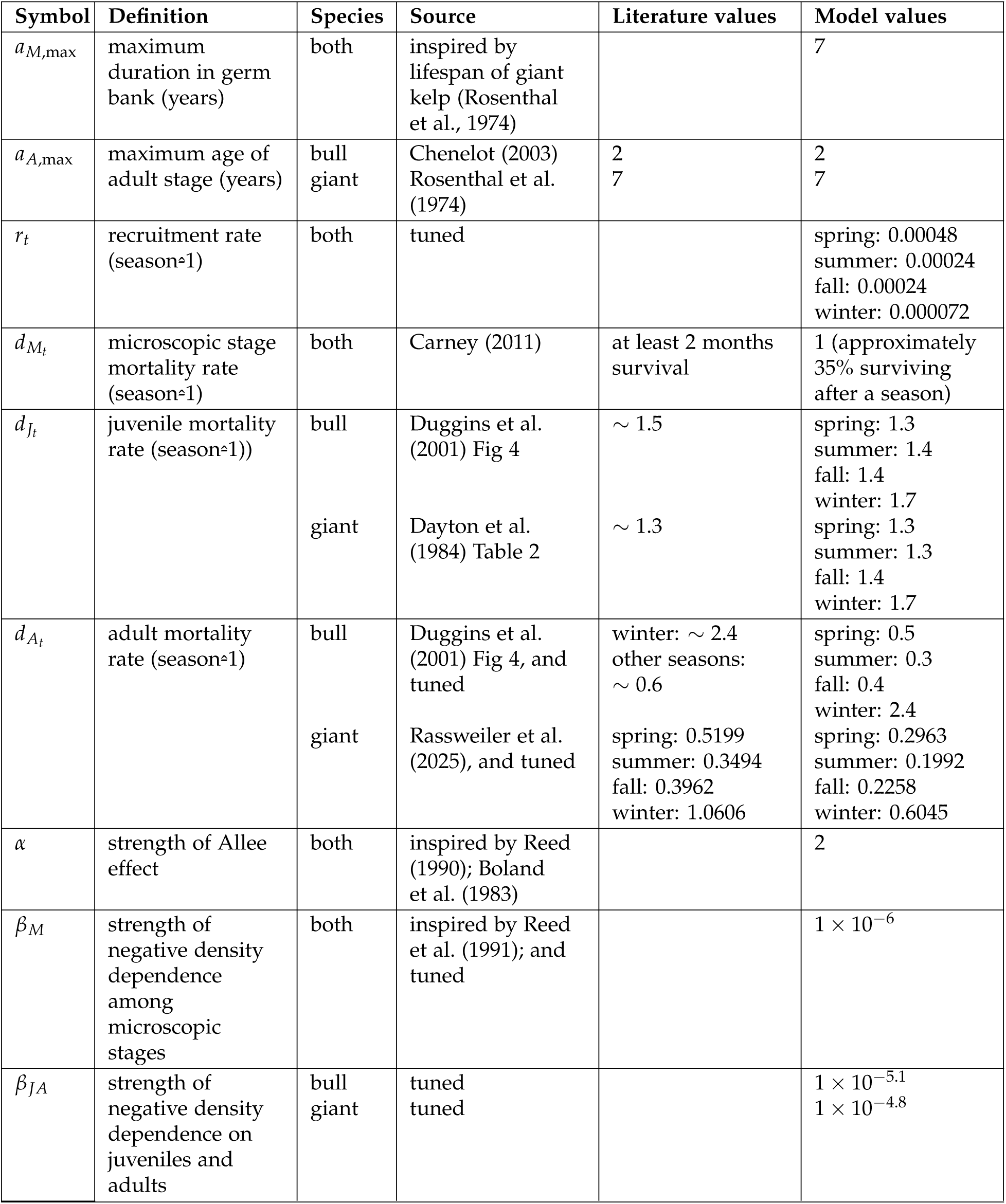
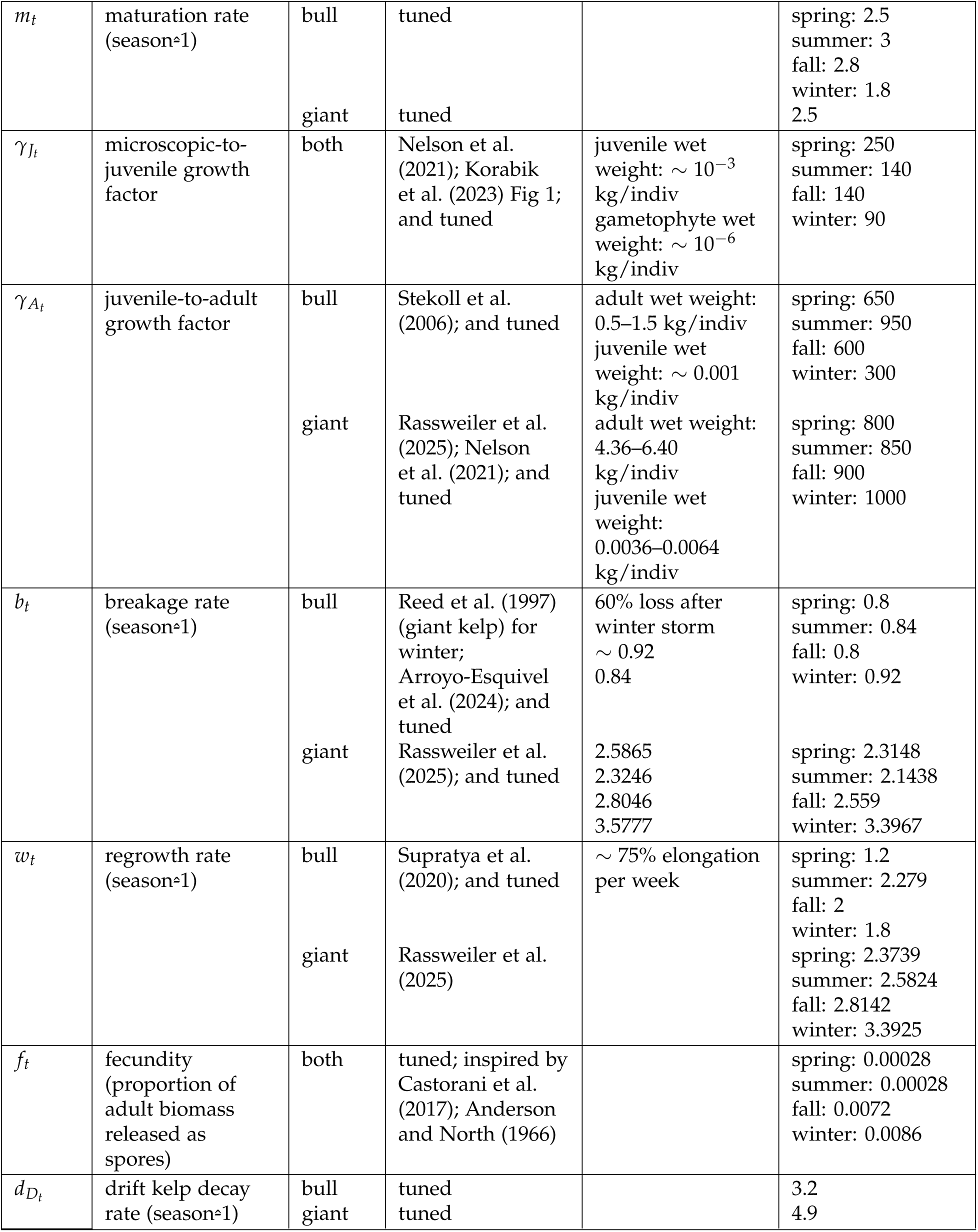
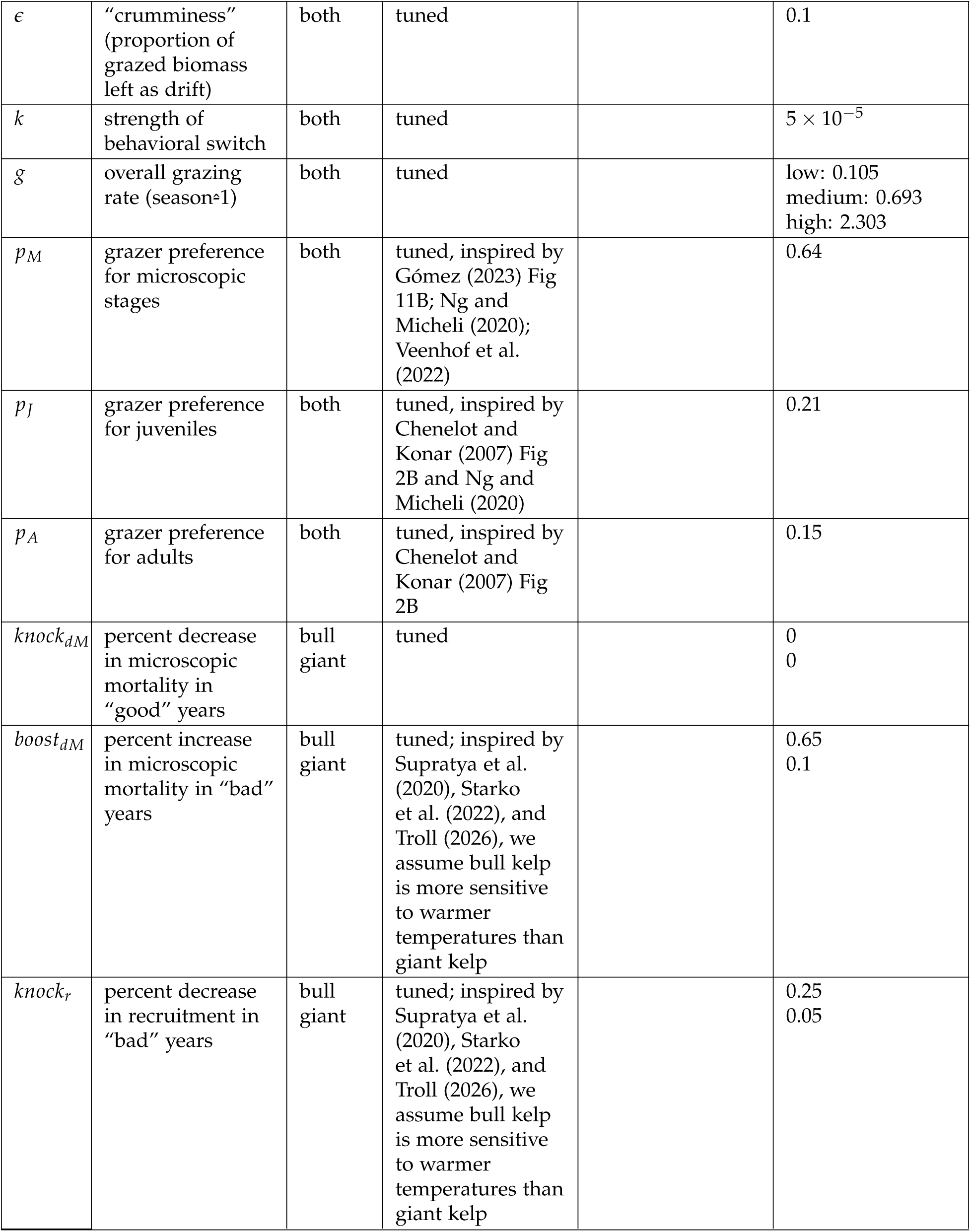
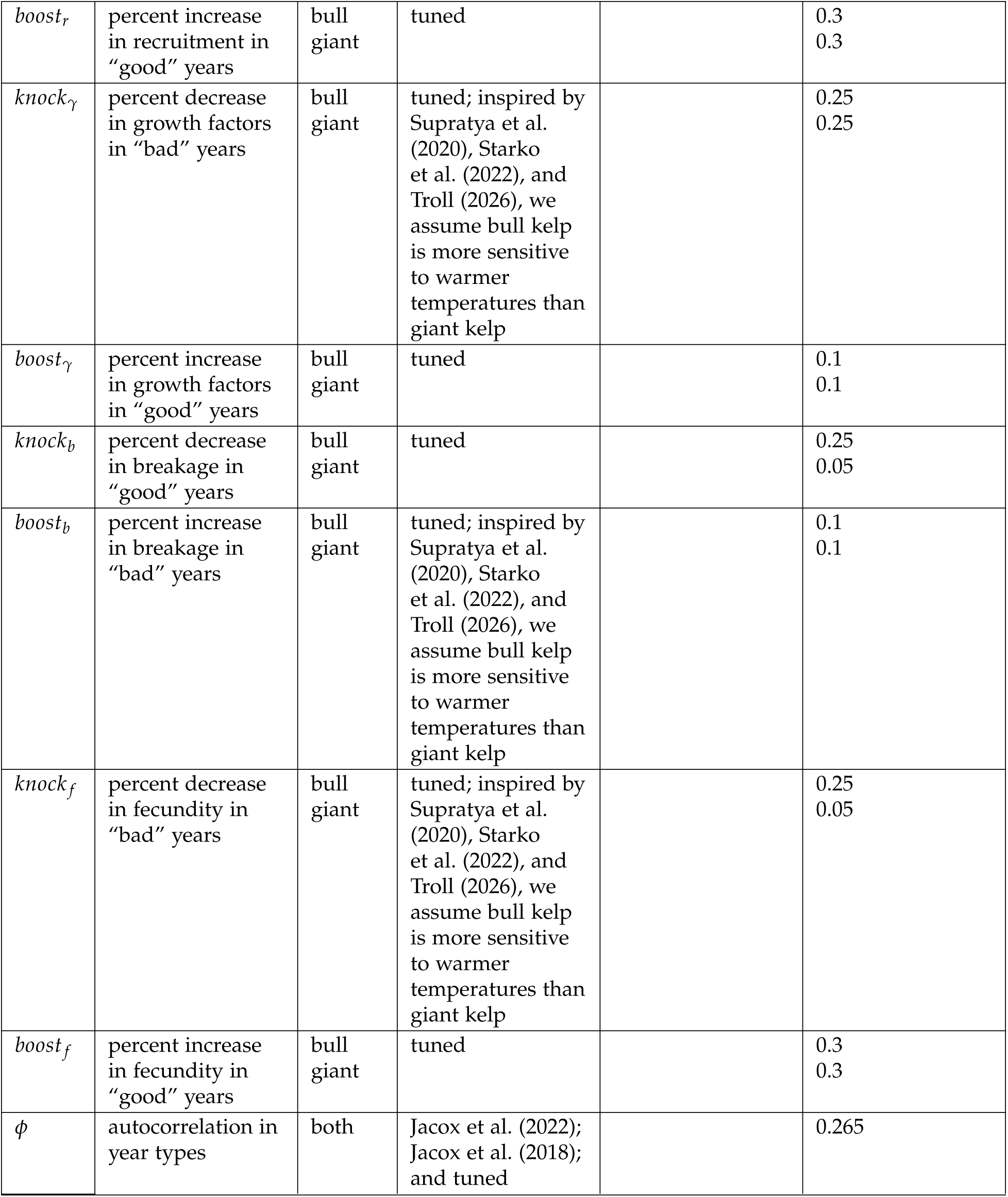
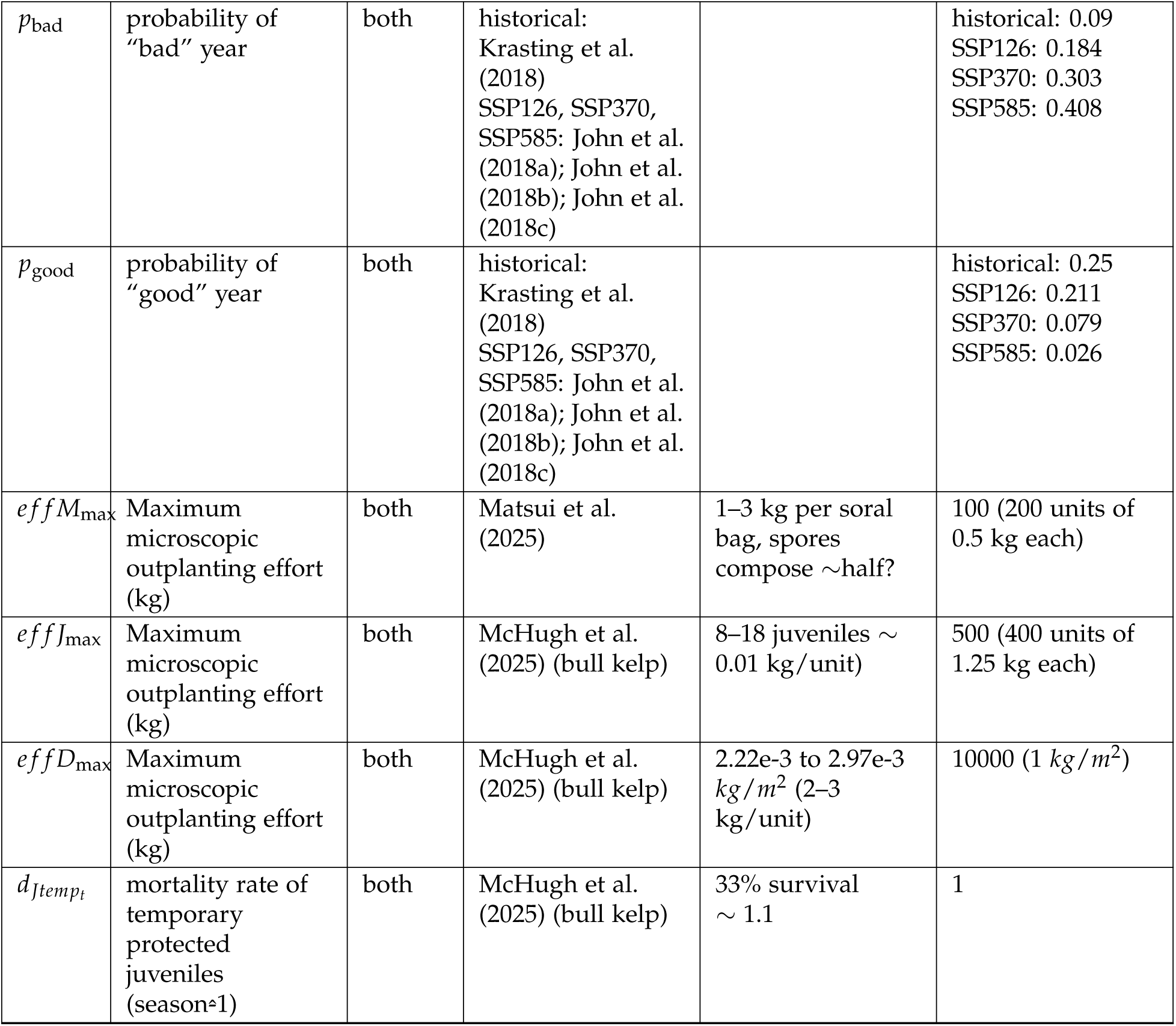
Model parameterization in the “neutral” year type.

| **Symbol** | **Definition** | **Species** | **Source** | **Literature values** | **Model values** |
|---|---|---|---|---|---|
| *a*<sub>*M*,max</sub> | maximum duration in germ bank (years) | both | inspired by lifespan of giant kelp (Rosenthal et al., 1974) |  | 7 |
| *a*<sub>*A*,max</sub> | maximum age of adult stage (years) | bull<br>giant | Chenelot (2003)<br>Rosenthal et al. (1974) | 2<br>7 | 2<br>7 |
| *r*<sub>*t*</sub> | recruitment rate (season≏1) | both | tuned |  | spring: 0.00048<br>summer: 0.00024<br>fall: 0.00024<br>winter: 0.000072 |
| *d*<sub>*M*<sub>*t*</sub></sub> | microscopic stage mortality rate (season≏1) | both | Carney (2011) | at least 2 months survival | 1 (approximately 35% surviving after a season) |
| *d*<sub>*J*<sub>*t*</sub></sub> | juvenile mortality rate (season≏1)) | bull<br>giant | Duggins et al. (2001) Fig 4<br>Dayton et al. (1984) Table 2 | ∼ 1.5<br>∼ 1.3 | spring: 1.3<br>summer: 1.4<br>fall: 1.4<br>winter: 1.7<br>spring: 1.3<br>summer: 1.3<br>fall: 1.4<br>winter: 1.7 |
| *d*<sub>*A*<sub>*t*</sub></sub> | adult mortality rate (season≏1) | bull<br>giant | Duggins et al. (2001) Fig 4, and tuned<br>Rassweiler et al. (2025), and tuned | winter: ∼ 2.4<br>other seasons: ∼ 0.6<br>spring: 0.5199<br>summer: 0.3494<br>fall: 0.3962<br>winter: 1.0606 | spring: 0.5<br>summer: 0.3<br>fall: 0.4<br>winter: 2.4<br>spring: 0.2963<br>summer: 0.1992<br>fall: 0.2258<br>winter: 0.6045 |
| *α* | strength of Allee effect | both | inspired by Reed (1990); Boland et al. (1983) |  | 2 |
| *β*<sub>*M*</sub> | strength of negative density dependence among microscopic stages | both | inspired by Reed et al. (1991); and tuned |  | 1 × 10<sup>−6</sup> |
| *β*<sub>*JA*</sub> | strength of negative density dependence on juveniles and adults | bull<br>giant | tuned<br>tuned |  | 1 × 10<sup>−5.1</sup><br>1 × 10<sup>−4.8</sup> |
| *m*<sub>*t*</sub> | maturation rate (season≏1) | bull<br>giant | tuned<br>tuned |  | spring: 2.5<br>summer: 3<br>fall: 2.8<br>winter: 1.8<br>2.5 |
| *γ*<sub>*J*<sub>*t*</sub></sub> | microscopic-to-juvenile growth factor | both | Nelson et al. (2021); Korabik et al. (2023) Fig 1; and tuned | juvenile wet weight: ∼ 10<sup>−3</sup> kg/indiv<br>gametophyte wet weight: ∼ 10<sup>−6</sup> kg/indiv | spring: 250<br>summer: 140<br>fall: 140<br>winter: 90 |
| *γ*<sub>*A*<sub>*t*</sub></sub> | juvenile-to-adult growth factor | bull<br>giant | Stekoll et al. (2006); and tuned<br>Rassweiler et al. (2025); Nelson et al. (2021); and tuned | adult wet weight: 0.5–1.5 kg/indiv<br>juvenile wet weight: ∼ 0.001 kg/indiv<br>adult wet weight: 4.36–6.40 kg/indiv<br>juvenile wet weight: 0.0036–0.0064 kg/indiv | spring: 650<br>summer: 950<br>fall: 600<br>winter: 300<br>spring: 800<br>summer: 850<br>fall: 900<br>winter: 1000 |
| *b*<sub>*t*</sub> | breakage rate (season≏1) | bull<br>giant | Reed et al. (1997) (giant kelp) for winter; Arroyo-Esquivel et al. (2024); and tuned<br>Rassweiler et al. (2025); and tuned | 60% loss after winter storm<br>∼ 0.92<br>0.84<br>2.5865<br>2.3246<br>2.8046<br>3.5777 | spring: 0.8<br>summer: 0.84<br>fall: 0.8<br>winter: 0.92<br>spring: 2.3148<br>summer: 2.1438<br>fall: 2.559<br>winter: 3.3967 |
| *w*<sub>*t*</sub> | regrowth rate (season≏1) | bull<br>giant | Supratya et al. (2020); and tuned<br>Rassweiler et al. (2025) | ∼ 75% elongation per week | spring: 1.2<br>summer: 2.279<br>fall: 2<br>winter: 1.8<br>spring: 2.3739<br>summer: 2.5824<br>fall: 2.8142<br>winter: 3.3925 |
| *f*<sub>*t*</sub> | fecundity (proportion of adult biomass released as spores) | both | tuned; inspired by Castorani et al. (2017); Anderson and North (1966) |  | spring: 0.00028<br>summer: 0.00028<br>fall: 0.0072<br>winter: 0.0086 |
| *d*<sub>*D*<sub>*t*</sub></sub> | drift kelp decay rate (season≏1) | bull<br>giant | tuned<br>tuned |  | 3.2<br>4.9 |
| *ϵ* | “crumminess” (proportion of grazed biomass left as drift) | both | tuned |  | 0.1 |
| *k* | strength of behavioral switch | both | tuned |  | 5 × 10<sup>−5</sup> |
| *g* | overall grazing rate (season≏1) | both | tuned |  | low: 0.105<br>medium: 0.693<br>high: 2.303 |
| *p*<sub>*M*</sub> | grazer preference for microscopic stages | both | tuned, inspired by Gómez (2023) Fig 11B; Ng and Micheli (2020); Veenhof et al. (2022) |  | 0.64 |
| *p*<sub>*J*</sub> | grazer preference for juveniles | both | tuned, inspired by Chenelot and Konar (2007) Fig 2B and Ng and Micheli (2020) |  | 0.21 |
| *p*<sub>*A*</sub> | grazer preference for adults | both | tuned, inspired by Chenelot and Konar (2007) Fig 2B |  | 0.15 |
| *knock*<sub>*dM*</sub> | percent decrease in microscopic mortality in “good” years | bull<br>giant | tuned |  | 0<br>0 |
| *boost*<sub>*dM*</sub> | percent increase in microscopic mortality in “bad” years | bull<br>giant | tuned; inspired by Supratya et al. (2020), Starko et al. (2022), and Troll (2026), we assume bull kelp is more sensitive to warmer temperatures than giant kelp |  | 0.65<br>0.1 |
| *knock*<sub>*r*</sub> | percent decrease in recruitment in “bad” years | bull<br>giant | tuned; inspired by Supratya et al. (2020), Starko et al. (2022), and Troll (2026), we assume bull kelp is more sensitive to warmer temperatures than giant kelp |  | 0.25<br>0.05 |
| *boost*<sub>*r*</sub> | percent increase in recruitment in “good” years | bull<br>giant | tuned |  | 0.3<br>0.3 |
| *knock*<sub>*γ*</sub> | percent decrease in growth factors in “bad” years | bull<br>giant | tuned; inspired by Supratya et al. (2020), Starko et al. (2022), and Troll (2026), we assume bull kelp is more sensitive to warmer temperatures than giant kelp |  | 0.25<br>0.25 |
| *boost*<sub>*γ*</sub> | percent increase in growth factors in “good” years | bull<br>giant | tuned |  | 0.1<br>0.1 |
| *knock*<sub>*b*</sub> | percent decrease in breakage in “good” years | bull<br>giant | tuned |  | 0.25<br>0.05 |
| *boost*<sub>*b*</sub> | percent increase in breakage in “bad” years | bull<br>giant | tuned; inspired by Supratya et al. (2020), Starko et al. (2022), and Troll (2026), we assume bull kelp is more sensitive to warmer temperatures than giant kelp |  | 0.1<br>0.1 |
| *knock*<sub>*f*</sub> | percent decrease in fecundity in “bad” years | bull<br>giant | tuned; inspired by Supratya et al. (2020), Starko et al. (2022), and Troll (2026), we assume bull kelp is more sensitive to warmer temperatures than giant kelp |  | 0.25<br>0.05 |
| *boost*<sub>*f*</sub> | percent increase in fecundity in “good” years | bull<br>giant | tuned |  | 0.3<br>0.3 |
| *ϕ* | autocorrelation in year types | both | Jacox et al. (2022); Jacox et al. (2018); and tuned |  | 0.265 |
| *p*<sub>bad</sub> | probability of “bad” year | both | historical: Krasting et al. (2018)<br>SSP126, SSP370, SSP585: John et al. (2018a); John et al. (2018b); John et al. (2018c) |  | historical: 0.09<br>SSP126: 0.184<br>SSP370: 0.303<br>SSP585: 0.408 |
| *p*<sub>good</sub> | probability of “good” year | both | historical: Krasting et al. (2018)<br>SSP126, SSP370, SSP585: John et al. (2018a); John et al. (2018b); John et al. (2018c) |  | historical: 0.25<br>SSP126: 0.211<br>SSP370: 0.079<br>SSP585: 0.026 |
| *effM*<sub>max</sub> | Maximum microscopic outplanting effort (kg) | both | Matsui et al. (2025) | 1–3 kg per soral bag, spores compose ∼half? | 100 (200 units of 0.5 kg each) |
| *effJ*<sub>max</sub> | Maximum microscopic outplanting effort (kg) | both | McHugh et al. (2025) (bull kelp) | 8–18 juveniles ∼ 0.01 kg/unit) | 500 (400 units of 1.25 kg each) |
| *effD*<sub>max</sub> | Maximum microscopic outplanting effort (kg) | both | McHugh et al. (2025) (bull kelp) | 2.22e-3 to 2.97e-3 *kg*/*m*<sup>2</sup> (2–3 kg/unit) | 10000 (1 *kg*/*m*<sup>2</sup>) |
| *d*<sub>*Jtemp*<sub>*t*</sub></sub> | mortality rate of temporary protected juveniles (season≏1) | both | McHugh et al. (2025) (bull kelp) | 33% survival<br>∼ 1.1 | 1 |

### Simulations and analysis

We wrote and compiled all code in R version 4.4.3. All simulations ran from an initial biomass of 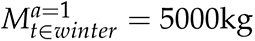 for 150 simulated years, and we conduct all interventions and analyses after the first 30 years of burn-in data are removed (Fig S3). We used the ggplot2 R package (Wickham, 2016) and Inkscape version 1.2 to generate and annotate figures. GPT-5.6 Luna assisted with coloring Fig 1.

For each simulated timeseries, we calculated five outcome metrics: First, summer adult biomass 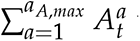 where *t* is the final summer of the simulation, as summer is when adult sporophyte biomass is at its maximum; Second, persistence as the proportion of years during which summer adult biomass 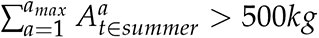 in 10,000*m*^2^. Third, fall microscopic biomass 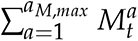 where *t* is the final fall of the simulation, as fall is when microscopic stage biomass is at its maximum; Fourth, grazing difference as the proportion of drift kelp biomass grazed in the summer minus the proportion of adult biomass grazed in the summer (1 *− e^−gD^t∈summer* ) *−* (1 *− e^−gA^t∈summer* ), averaged across summers in the simulation; Lastly, we collect the number years during which intervention occurred in the simulation.

To quantify the relative influence of intervention decisions and biophysical attributes on outcome metrics, we conducted a global sensitivity analysis following an algorithm similar to Harper et al. (2011). First, we set the species, grazing, and climate context for which to run simulations (Table 1). Next, we drew 1000 random samples for the intervention decisions: cue, threshold, wait time, effort, and life stage at introduction. For each of the 1000 sets of intervention decisions, we ran 100 stochastic simulations and saved the averages of their outcome metrics alongside the corresponding intervention and biophysical context information into an R dataframe.

We used the randomForest R package (Liaw and Wiener, 2002) to run regression of each outcome metric on the intervention and biophysical variables, and rank the importance of those variables for each outcome metric (Fig 5). Importance of a variable is the percent decrease in MSE of the predicted outcome value when the variable is removed from randomForest’s regression model.

**Figure 5:**
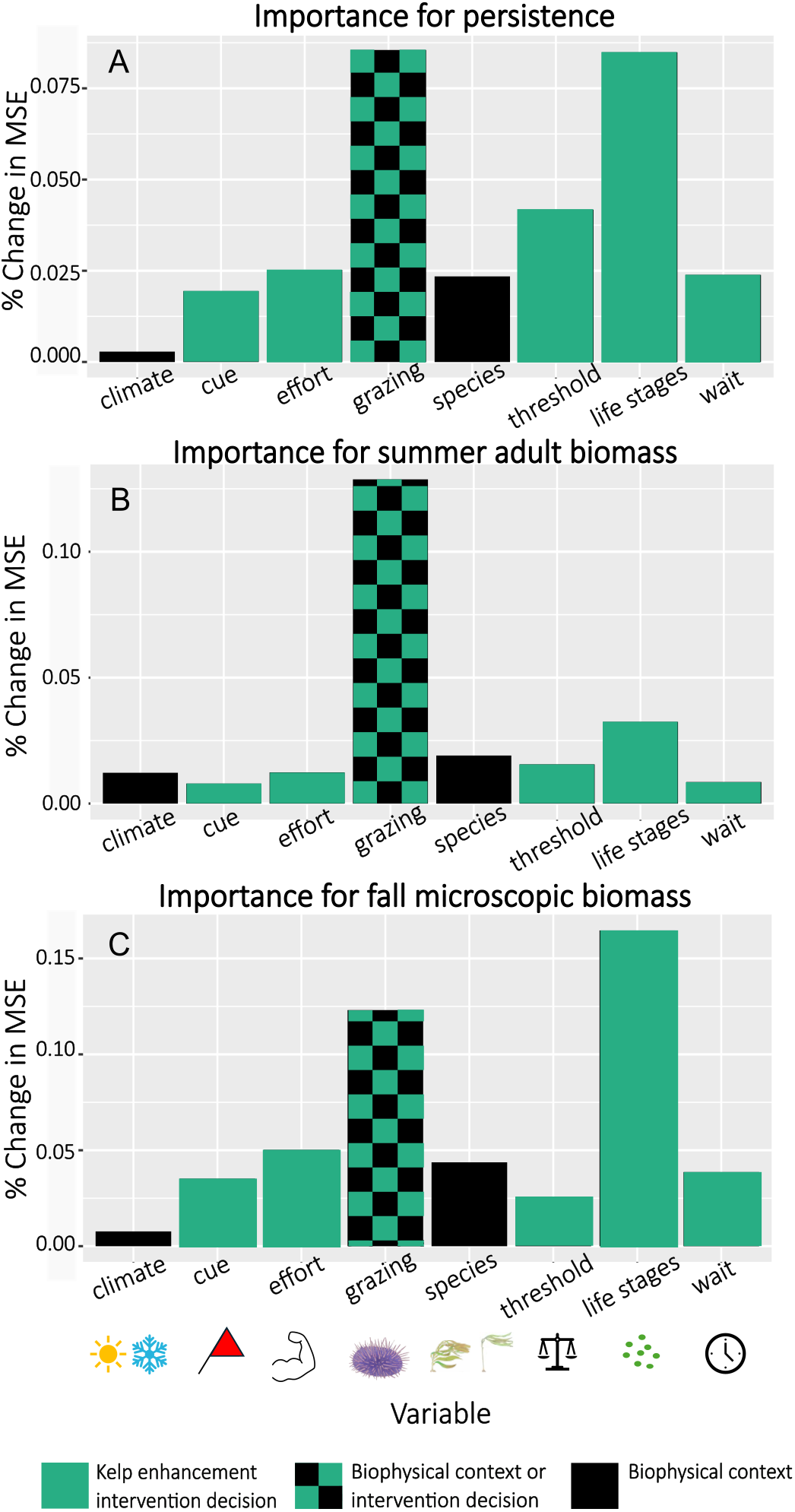
Relative importance of intervention decisions and biophysical context. Importance of variables for the following outcome metrics: A) population persistence, B) summer adult biomass, C) fall microscopic biomass. Importance is measured as the % decrease in MSE when the respective variable is removed from a random forest regression model predicting the outcome metric. Teal-green and black checkerboard labels grazing intensity, a variable that is both an intervention decision and a feature of the surrounding biophysical context.

To compare intervention efficacy between different life stages at introduction, we partitioned the above-described simulation data by species, grazing scenario, and climate scenario, and plotted the distributions of each outcome metric resulting from each set of introduced life stages (Fig 6).

**Figure 6:**
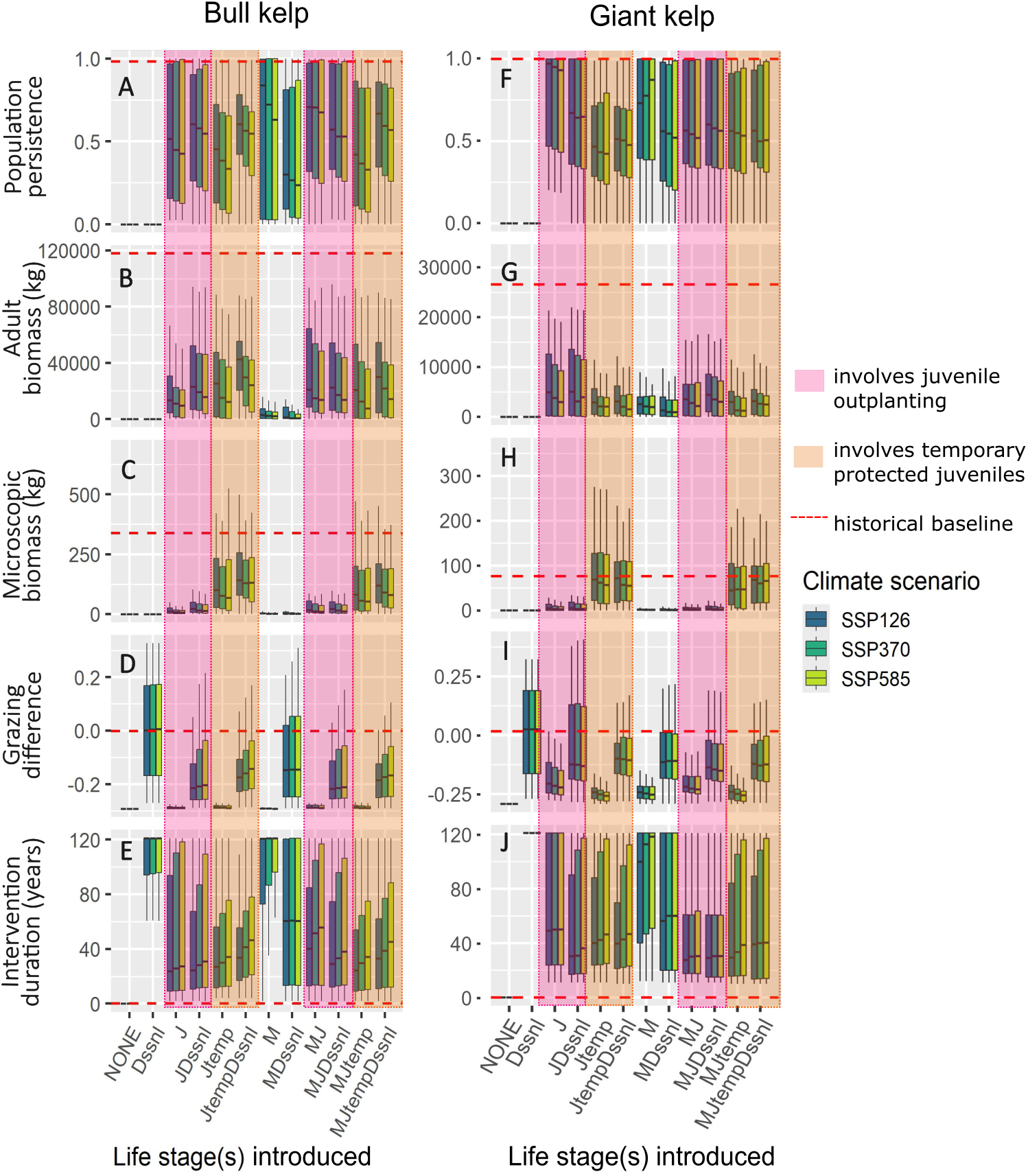
Effects of life stages at introduction. Box plots showing distributions of outcome metrics across sets of introduced life stages (horizontal axes), climate scenario (color), and species (columns). Results are from simulations under “high” grazing. Outcome metrics: A) and F) population persistence; B) and G) summer adult biomass; C) and H) fall microscopic biomass; D) and I) proportion of drift biomass grazed minus proportion of adult biomass grazed; E) and J) intervention duration. Each box plot, except for those of type ”NONE”, summarizes 90 data points. Each data point represents the mean of 100 stochastic simulations of a particular intervention case, i.e., combination of intervention cue, threshold, effort, type, wait time, and climate scenario. Dotted red lines indicate the mean of the respective outcome metric across 100 stochastic simulations in the “historical” climate scenario without intervention. Shaded pink rectangles (left) highlight interventions that use juvenile outplants. Shaded orange rectangles (right) highlight intervention types that use temporary protected juveniles.

To understand how wait time relates to intervention outcomes and total intervention effort, we used the partitioned simulation data to plot outcome metrics against the product of intervention duration and intervention effort used in the corresponding simulations, and also colored the points based on the wait time used in those simulations (Fig 7,8).

**Figure 7:**
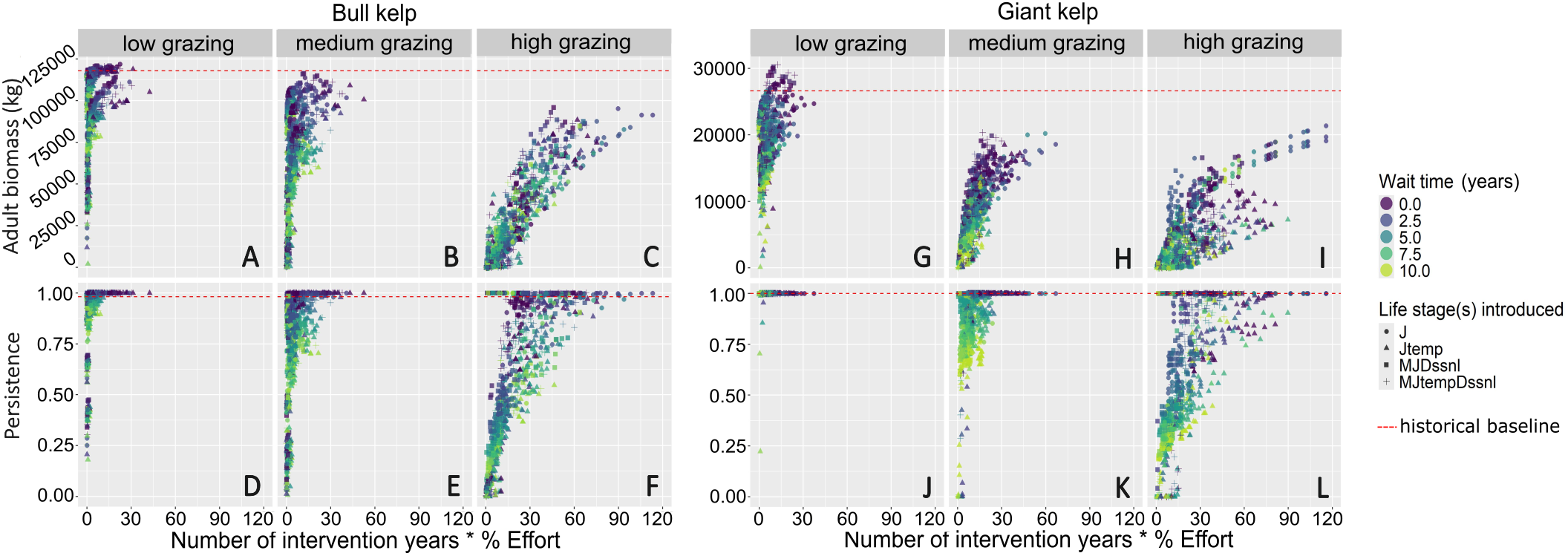
Effects of intervention effort and wait time. Summer adult biomass (A-C, G-I) and population persistence (D-F, J-L) plotted against total intervention effort, across “low” (A, D, G, J), “medium” (B, E, H, K) and “high” (C, F, I, L) grazing scenarios. Bull kelp intervention outcomes (A-F). Giant kelp intervention outcomes (G-L). The maximum value on the horizontal axis indicates 120 years of 100% effort. Each point represents the mean of 100 stochastic simulations of a particular intervention case, i.e., combination of intervention cue, threshold, effort, type, wait time, and grazing and climate scenarios. Dotted red lines indicate the mean across 100 stochastic simulations of each outcome metric in the “historical” climate scenario without intervention. Points are colored by wait time and shaped by intervention type.

**Figure 8:**
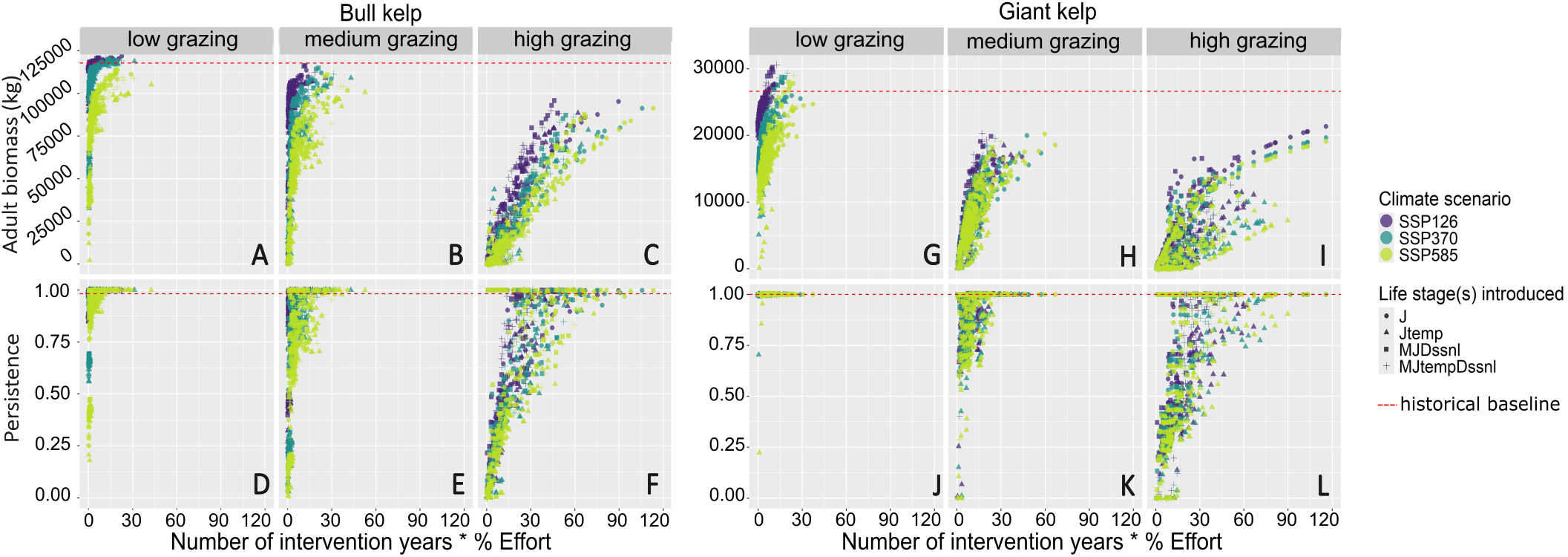
Effects of intervention effort and climate. Summer adult biomass (A-C, G-I) and population persistence (D-F, J-L) plotted against total intervention effort, across “low” (A, D, G, J), “medium” (B, E, H, K) and “high” (C, F, I, L) grazing scenarios. Bull kelp intervention outcomes (A-F). Giant kelp intervention outcomes (G-L). The maximum value on the horizontal axis indicates 120 years of 100% effort. Each point represents the mean of 100 stochastic simulations of a particular intervention case, i.e., combination of intervention cue, threshold, effort, type, wait time, and grazing and climate scenarios. Dotted red lines indicate the mean across 100 stochastic simulations of each outcome metric in the “historical” climate scenario without intervention. Points are colored by climate scenario and shaped by intervention type.

## Results

### Model Behavior

Simulated adult biomass for bull and giant kelp in the “historical” climate scenario (Fig 4A) roughly align with the empirical estimates from Landsat (Fig 1). Bull kelp biomass is on the order of 10^5^*kg* while giant kelp biomass is on the order of 10^4^*kg*. For both species, there is about one peak in biomass per year (summer).

In our model simulations, intervention improves population persistence and biomass outcomes in the “moderate” climate scenario (SSP370). With intervention, bull and giant kelp persist at a biomass range comparable to that of the “historical” climate scenario (Fig 4C) when the populations would otherwise perish without intervention (Fig 4B). The average biomass for giant kelp under “moderate” climate with intervention is about 25% lower than the “historical” scenario without intervention.

### Grazing and life stages at introduction are the most influential intervention decisions for population outcomes

In our global sensitivity analysis of variables that determine biophysical context and intervention approach, we find that population persistence and summer adult biomass are most sensitive to grazing intensity and life stages at introduction (Fig 5A,B).

When looking at their interaction, we see that the negative effect of grazing on population persistence is strongest when temporary protected juveniles are introduced, and the negative effect of grazing on adult biomass is strongest when microscopic outplants are introduced (Fig S4). Grazing intensity reflects both an intervention decision (i.e., grazer removal, predator reintroduction) and the surrounding biophysical context (e.g., naturally occurring grazer abundance/activity, presence of predators, environmental conditions).

Sensitivity to life stages at introduction indicates that the decision of which combination of life stages to use for restoration is more influential than most other aspects of the intervention decision space. Microscopic biomass (i.e., the germ bank) responds more strongly to life stages at introduction than any other intervention decision or biophysical variable that we tested (Fig 5C). This leads us to a possible mechanism for effective kelp enhancement interventions: adding to the germ bank.

### Temporary protected juveniles restore germ banks, but the need for germ bank restoration differs between species

In our comparison of kelp enhancement approaches, we find that the optimal life stages at introduction differ between species. Of the combinations of life stages to introduce, those that combine temporary protected juveniles with drift kelp supplementation, i.e., “JtempDssnl” and “MJtempDssnl” lead to the greatest median adult biomass for bull kelp (orange highlights in Fig 6B). In contrast, the interventions that maximize adult giant kelp biomass are those that involve juvenile outplanting: “J”, “JDssnl”, “MJ”, and “MJDssnl” (pink highlights in Fig 6G). These are also the interventions that lead to the greatest persistence for both species while recovering biomass (pink highlights in Fig 6A,F).

For both species, interventions that involve temporary protected juveniles lead to the greatest recovery of germ bank biomass compared to other strategies (orange highlights in Fig 6C,H). These interventions, i.e., “Jtemp”, “JtempDssnl”, “MJtemp”, and “MJtempDssnl”, also reduce total intervention duration compared to other strategies for bull kelp (orange highlights in Fig 6E), suggesting that germ bank recovery via temporary protected juveniles is relatively self-sustaining and does not require continuous intervention effort for bull kelp.

Although strategies that involve temporary protected juveniles restore the “historical” germ bank biomass of giant kelp (orange highlights in Fig 6H), the benefits of these strategies do not extend to giant kelp adult biomass, persistence, or intervention duration compared to interventions that involve juvenile outplanting.

Despite contributing directly to the germ banks, microscopic outplants alone (“M”) do not restore biomass of either species (Fig 6B,C,G,H) and microscopic outplanting effort is nearly continuous (i.e. intervention duration; Fig 6E,J) in order to achieve persistence (Fig 6A,F). Seasonal drift kelp supplementation alone (“Dssnl”) also does not restore biomass and persistence for either species (Fig 6A-C,F-H), despite shifting grazing pressure away from live kelp to resemble “historical” grazing behavior (Fig 6D,I).

The differences between intervention strategies are more pronounced with increasing grazing intensity (Fig S4). While variance in intervention outcomes is primarily driven by climate scenario under “low” grazing, differences between sets of introduced life stages is the dominant source of variance under “high” grazing.

### Waiting on kelp enhancement can save effort but sacrifices outcomes

In our analysis of intervention effort and wait time, we find that waiting longer before starting intervention reduces biomass and persistence outcomes while also reducing overall intervention effort. Across all panels, biomass and persistence outcomes increase and eventually saturate with increasing total effort. Intervention approaches which used longer wait time (lighter colors) generally fall in the lower left regions of the distributions, where adult biomass, persistence, and total intervention effort are all reduced. In contrast, intervention approaches which used shorter wait time (darker colors) lead to higher biomass and persistence outcomes through greater and more frequent effort.

When comparing between species, we find that the effect of wait time on intervention outcomes and total effort is stronger for giant kelp than it is for bull kelp.

Whereas there is a clearer gradient between short and long wait times along the low-effort-low-outcome to high-effort-high-outcome axis for giant kelp (Fig 7G-L), there appear to be more cases for bull kelp in which rapid intervention (darker colors) leads to similar outcomes and total effort as does waiting longer (lighter colors).

Grazing intensity also plays a role in the effectiveness of kelp enhancement interventions (consistent with Fig 6). Under “low” grazing intensity, interventions meet or exceed historical baselines for both species when they start within three years of their intervention thresholds being met (Fig 7A,D,G,J). Full recovery of historical baselines requires less than fifteen years of maximum kelp enhancement effort. As grazing intensifies, less recovery of adult biomass and persistence is possible and greater overall intervention effort is needed to achieve comparable biomass recovery as in the “low” grazing scenario (see maximum outcome values in Fig 7B,E,H,K and C,F,I,L compared to A,D,G,J, respectively). Under “high” grazing, even maximum intervention effort and immediate intervention does not recover “historical” biomass, although persistence is possible. The cases with greatest biomass recovery tend to occur with long-term high intervention effort and wait times under five years.

Climate scenario also limits recovery potential. When we color the same plots by climate instead of wait time, we find that more intense climate scenarios lead to lower distributions for adult biomass and persistence outcomes, while the relationships between intervention effort and outcomes are consistent across climate scenarios (Fig 8).

## Discussion

### Implications for kelp restoration

Our results suggest that a dual intervention approach is effective for kelp restoration: reduce grazing to protect kelp and employ kelp enhancement to rebuild population size. This aligns with results from pilot restoration projects in California and abroad (Eger et al., 2022; Ward et al., 2022; public communication from Giant Giant Kelp Restoration Project), and other modeling work (Arroyo-Esquivel et al., 2023; White et al., 2026 and personal communication with Kim et al., *in review*). High sensitivity of adult biomass to grazing intensity is consistent with our understanding of kelp forest and urchin barren alternative stable state dynamics: the behavioral switch from passive to active grazing drives kelp collapse (Arroyo-Esquivel et al., 2023; Hopf et al., 2022; Steneck and Johnson, 2013). Grazer suppression can be achieved via direct removal, predator reintroduction, or short-term deterrence with drift kelp supplementation. Our results suggest that outplanting juveniles directly to the reef can be an effective strategy for giant kelp restoration, while protecting the juvenile outplants (e.g., by elevating them off the benthos using an ARKEV-like structure) for the purpose of reseeding a kelp bed can be especially helpful for bull kelp.

A potentially surprising result is the relative inefficacy of drift kelp supplementation under our model parameterization, even despite shifting grazing away from live kelp. We suspect that this reflects steep hysteresis and suggests either that much more and/or frequent drift kelp supplementation than the maximum effort we simulated (1 *kg*/*m*^2^ seasonally) would shift the system out of an urchin barren state, or that drift kelp supplementation is ultimately unlikely on its own to shift the system without additional kelp enhancement. In a pilot restoration study, McHugh et al. (2025) saw that drift kelp supplementation of 2.22e-3 to 2.97e-3 *kg*/*m*^2^ significantly reduced grazing on their assays in three sites after 24 hours. Taken together, we affirm that drift kelp supplementation is a promising tool for short-term deterrence of grazing at targeted sites to protect new outplants, but implementation at a scale necessary to satiate grazers and shift their behavior across a population would require frequent and abundant resupply from farm or lab based growers.

Microscopic outplants alone also could not restore adult biomass of either species, despite contributing directly to the germ bank, consistent with the challenges that restoration projects using green gravel, seeded twine, or soral bag techniques face in outplanting enough surviving microscopic material to match natural fecundity (Graham et al., 2023; Reed et al., 2024). When we increased microscopic outplanting effort from 100*kg* to 500*kg* in our model, intervention outcomes improved, confirming that microscopic outplanting success is limited by scale. Using temporary protected juveniles helps address the issue of scale by reseeding the kelp bed in amounts difficult to achieve by direct microscopic outplanting (McHugh et al., 2025). The maximum effort for temporary protected juveniles that we used in our model was 500*kg*, which represents an optimistic scaling to 1 ARKEV unit per 25*m*^2^, each carrying 1.25 *kg* juvenile sporophytes, compared to McHugh et al.’s (2025) trial of roughly 0.05 units per 25*m*^2^ at 0.01 *kg* per unit. This would require abundant but not unrealistic supply of juvenile sporophytes from current growers in California.

Juvenile sporophytes outplanted directly to the environment were effective for giant kelp restoration, while bull kelp benefited most from the temporary protected juveniles. In addition to their differential reliance on germ banks (see subsection *Broader insights for highly variable systems*), this result might be due to higher juvenile survival for giant kelp than bull kelp in our parameterization. The newly developed ”holdfast-graft” method for outplanting juvenile kelp sporophytes could help improve survival and scaling of juvenile kelp outplants for both species in California (Ling et al., 2026). The method, developed in Tasmania for giant kelp restoration, involves tying short 60*cm* lengths of seeded twine onto naturally occurring holdfast stubs of other macroalgal species in the field. This method takes a diver 45 seconds per holdfast and improves reef attachment by 40-fold compared to previous juvenile outplanting methods via seeded twine tied to rocks and juvenile sporophytes grown on stones (Ling et al., 2026). We estimate this method would take at least 19 hours of underwater dive time to attach our model’s maximum effort of 500 *kg* juvenile outplants, not counting the costs and time it would take to locate and move to suitable holdfasts across the kelp bed.

We found that interventions in a naturally fluctuating kelp system were most effective when initiated sooner, i.e., within about five years of persistent decline. In other kelp restoration models, Hopf et al. (2025) and White et al. (2026) show that early, even preemptive, intervention optimizes outcomes. Here, we emphasize that rapid intervention can in some cases prove unnecessary. Under “low” and “medium” grazing in our model, it was possible to meet or exceed “historical” baselines when intervening within three years. In a real world example on Wheeler North Reef in Southern California, natural giant kelp recruitment from a nearby source population exceeded the density of outplanted lab grown recruits by 75-fold, highlighting the immediate relevance of wait time to feasibility and cost of kelp enhancement (Reed et al., 2024).

While threshold of decline was less influential than other intervention decisions in our analyses, we find that using a percent rather than percentile threshold leads to more consistent intervention across species. This happens because percentiles are sensitive to the width of biomass distributions, which differ between our two species. Our modeled biomass for giant kelp was less variable across simulations than bull kelp, which means that for the same percentile threshold, intervention was triggered more often in giant kelp than bull kelp. A percent threshold would trigger intervention at the same relative decline across species. When we tested a percent rather than percentile threshold, species identity became less influential on outcomes (Fig S5). Threshold was not highly ranked in importance for biomass, whether defined by percent or percentile, but managers might find it helpful to consider the species-sensitivity of these metrics.

### Caveats and assumptions

Our model, as with any model, inevitably makes simplifying assumptions.

Regarding grazing dynamics, we assume that grazer behavior rather than abundance drives kelp population dynamics and we assume with high uncertainty a value for the behavioral switch parameter *k*. By making the first assumption, we omit a feedback between kelp and grazer abundance. However, if urchin density is driven primarily by recruitment (Hereu et al., 2012; Ling et al., 2019) and urchin recruitment is not strongly related to kelp abundance (Baskett and Salomon, 2010; Hereu et al., 2012), then missing this link should not affect the realism of our model much. If grazer abundance were dynamically linked to kelp in our model, we might expect kelp enhancement interventions to be less effective than we see under constant overall grazing intensity, as grazer abundance would increase whenever kelp biomass increases. Under our second assumption, about equal proportions of drift and adult kelp were consumed each season in the baseline condition. We doubt if this realistically reflects passive urchin grazing behavior, which primarily concentrates on drift kelp (Harrold and Reed, 1985; Kriegisch et al., 2019). If grazing behavior in our model was more concentrated on available drift kelp (higher *k*), we would expect greater probability of natural recovery and less intervention effort used to restore “historical” baselines. We highlight the need for more concrete field data on stage-specific grazing rates and their sensitivities to kelp abundance to better resolve kelp forest grazing dynamics.

Most biomass in the adult stages of either species did not survive longer than one year in our model (see Fig S6). Given that the lifespan of giant kelp can be up to seven years in wave-sheltered locations, we expected its adult age structure to be deeper (i.e., older, longer-lived) in our model (Rosenthal et al., 1974). We suspect that a deeper age structure for giant kelp would enhance the difference we see in the species’ reliance on a germ bank for recovery. Following from White et al. (2022), we also expect that a deeper age structure would slow the rate of natural recovery to enable longer intervention wait times for giant kelp. Age structured data for kelp, while challenging to obtain because it entails tracking individual sporophytes through fast biomass turnover, would improve the ability to parameterize models like this and resolve optimal restoration approaches.

Intervention threshold and wait time are correlated in their influences on outcome metrics. Due to their correlation, we risk understating the influence of threshold as we focus our analysis on wait time in this paper. The decisions to initiate intervention at a higher biomass threshold or with shorter wait time both independently lead to more frequent intervention and thus greater number of intervention years (compare Fig 7 with Fig S7). Some variance in intervention outcomes can thus be explained by both threshold and wait time, which might explain why these variables rank similarly in importance. However, we confirmed that wait time indeed correlates more strongly with both adult biomass and persistence outcomes than does threshold (Fig S8, note exception of stronger correlation with threshold in the low grazing case for giant kelp). Future modeling and empirical work could further explore tradeoffs associated with delayed intervention and identify the conditions under which either the threshold of decline or duration of decline are influential for intervention outcomes.

### Broader insights for highly variable systems

Our results contribute to a body of work investigating population buffering by germ banking in highly variable systems (Childs et al., 2010; Cohen, 1966; Edwards, 2022; Evans and Dennehy, 2005; Hoffmann and Santelices, 1991). Our finding that the annual bull kelp benefited most from interventions which enhanced its germ bank, while the perennial giant kelp could recover more adult biomass without such interventions suggests that organisms with life histories that provide redundant buffering in later life stages (e.g., flexibility to distribute reproduction over time) might not rely as much on buffering from a dormant germ stage. Early models investigating germ banking and adult longevity showed that the two strategies provide analogous and potentially redundant demographic buffering amid environmental variability (Rees, 1993, 1994; Tuljapurkar and Wiener, 2000). This led to the hypothesis that annual and semelparous species are more sensitive to disruptions to their germ bank than are perennial and iteroparous species, because the latter benefit from multiple cohorts and future opportunities to both germinate and reproduce, while the former have access only to the germ bank and current reproductive cohort (reviewed in Friedman, 2020).

While some empirical evidence from plant taxa shows that seed dormancy accounts for annual species’ recovery and persistence through fire, drought, and temperature variability (Larios et al., 2013; Plue et al., 2021), and contributes less to population outcomes for perennials (Evans et al., 2007; Larios et al., 2013; Thompson et al., 1998), the literature so far is not conclusive on whether semelparous species in general rely on their germ banks more than their iteroparous counterparts (Evans et al., 2007). Our results for bull and giant kelp provide support for this hypothesis from a system outside the usual terrestrial desert and grassland plants of this life history literature.

Life history also influences a population’s natural recovery time, which affects how long managers might be willing to “wait-and-see”. Longer-lived species recover slower from disturbance than shorter-lived species, as it takes time for demographic effects to propagate through the population’s age structure (Capdevila et al., 2022; White et al., 2022). For longer-lived iteroperous species compared to shorter-lived semelparous species, we might expect slower natural recovery from disturbance and slower response to intervention. Drawing on literature that highlights the significant role that germ banks play in the natural recovery of annual species (Larios et al., 2013; Plue et al., 2021), we might also expect that if the germ stage is itself long-lived and forms an age-structured bank, then annual populations could also naturally recover over longer timescales, meaning the potential to wait longer on intervention. For species, like bull kelp, that lack conclusive evidence for a long-lived germ bank, we highlight the need to determine germ bank longevity. Across highly variable systems, we encourage managers and researchers to identify intervention strategies which lead to rapid responses in population outcomes, and investigate how environmental variability and species’ life history figure into those relationships.

While delaying intervention to wait for natural recovery or better information can save costs and effort, the “wait-and-see” approach has potentially nuanced social implications. Depending on the context, resource-dependent livelihoods and important cultural practices might be at stake, and managers’ decision to not intervene or to ask for public patience can lead to conflict, loss of trust, and perception of institutional neglect for a community’s interests and priorities (Bach et al., 2022; Lö fqvist et al., 2023). In one example, delayed response to reef fish population declines in the 1990’s in the Gulf of Mexico vermillion snapper fishery eroded fishers’ trust in the Gulf of Mexico Fishery Management Council (Gulf Council) and NOAA Fisheries, and frustration among fishers persists to the 2020’s (public communication by Guindon, 2022). The Gulf Council eventually restricted commercial fishing in 2005, claiming that additional data was needed to make the decision (Council, 2004; Service, 2005).

In this paper, we made the critical assumption that the goal of intervention is to restore biomass to a historical state. However, there exists a wide range of possible intervention goals that address diverse community priorities, such as deeply-held preferences and attitudes around intervention (Enevoldsen, 2024) and priorities around cultural stewardship and use (Lyver et al., 2016; Wehi and Lord, 2017). Additionally, especially in highly variable systems that might not have a well-understood historical state, a return to past baselines might not be feasible (Hobbs and Harris, 2001). In such cases, the Resist-Accept-Direct (Schuurman et al., 2020) and related adaptive management (Millar et al., 2007) frameworks become relevant for flexible, socially responsible, and future-minded restoration.

## Acknowledgments

MZB acknowledges support from the National Science Foundation Graduate Research Fellowship Program #2439024. MLB and MZB acknowledge support from the National Science Foundation Dynamics of Integrated Socio-Environmental Systems #2108002. MLB acknowledges support from University of California Agricultural Experiment Station Project #CA-D-ESP-2244-H. MZB acknowledges assistance from GPT-5.6 Luna in coloring Fig 1 and searching the literature. The authors thank Dr. Jay Stachowicz for his friendly review of the original draft, and Calvin Munson, Kyumin Kim, and Drs. Jorge Arroyo-Esquivel, Nicholas Burnett, Katie Dobkowski, and Mark Carr for their comments and suggestions during the conception of the model. The authors are grateful to Drs. Andrew Rassweiler and Dan Reed for their clarifying insights on giant kelp data from the SBC LTER dataset, and Dr. Lauran Liggan for her insights on likely *Nereocytstis* gametophyte abundances. The authors also thank Tristin McHugh, Bennet Bugbee, and Dr. Scott Hamilton for discussions on current and ongoing kelp restoration approaches.

## Author contributions

MZB conceived the model, developed the code, and wrote the original draft. MLB advised and oversaw project progress. All authors reviewed and edited the writing.

