## Supplemental Text for "Restoration decisions in highly variable systems: Whether, when, and how to enhance kelp"

### Appendix S1:

### Restoration decisions in ephemeral systems: Whether, when, and how to enhance kelp

Mei Z. Blundell[1,*]

Marissa L. Baskett [1]

1. Department of Environmental Science and Policy, University of California Davis, Davis, California, USA;

### Additional Methods

#### *Matrix model*

We consolidate kelp demographic dynamics into a matrix of transitions between all stages and ages. The biomass distribution at the beginning of each season $t$ is

$$\vec{X}_t = \begin{pmatrix} M_t^1 \\ \vdots \\ M_t^7 \\ J_t \\ J_{temp_t} \\ A_{temp_t} \\ A_t^1 \\ \vdots \\ A_t^7 \\ D_t \end{pmatrix} \tag{S1}$$

where superscripts $a$ for the $M_t^a$ and $A_t^a$ entries represent age, i.e., number of years that biomass has remained in the microscopic or adult life stage.

Using the transition terms described in *Demographic dynamics* in the main *Methods*, $\vec{X}_t$ is updated to the next season according to

$$\vec{X_{t+1}} = A_t \vec{X}_t \tag{S2}$$

given the transition matrix

$$A_{t \notin winter} = \begin{bmatrix} S_{M_t^1} & & & & & F_t^1 & F_t^1 & \cdots & F_t^7 & \\ & \ddots & & & & & & & & \\ & & S_{M_t^7} & & & & & & & \\ R_t^1 & \cdots & R_t^7 & S_{J_t} & & & & & & \\ & & & & S_{J_{temp_t}} & & & & & \\ & & & & G_{temp_t} & S_{A_{temp_t}} & & & & \\ & & & G_t & & & S_{A_t^1} & & & \\ & & & & & & & \ddots & & \\ & & & & & & & & S_{A_t^7} & \\ & & & & & B_{temp_t} & B_t^1 & \cdots & B_t^7 & S_{D_t} \end{bmatrix} \tag{S3}$$

for all transitions between seasons other than winter-to-spring. The winter-to-spring transition differs because it accounts for "aging up" the microscopic and adult stages by one year. We do this by shifting the survival terms $S_{M_t^a}$ and $S_{A_t^a}$ down to the subdiagonal:

$$A_{t \in winter} = \begin{bmatrix} & & & & & F_t^1 & F_t^1 & \cdots & F_t^7 & \\ S_{M_t^1} & & & & & & & & & \\ & \ddots & S_{M_t^6} & & & & & & & \\ R_t^1 & \cdots & R_t^7 & S_{J_t} & & & & & & \\ & & & & S_{J_{temp_t}} & & & & & \\ & & & & G_{temp_t} & S_{A_{temp_t}} & & & & \\ & & & G_t & & & & & & \\ & & & & & & S_{A_t^1} & & & \\ & & & & & & & \ddots & S_{A_t^6} & \\ & & & & & B_{temp_t} & B_t^1 & \cdots & B_t^7 & S_{D_t} \end{bmatrix}. \tag{S4}$$

We make sure that adult biomass does not exceed the species-specific maximum age by setting $S_{A_t}^a = 0$ when $a > a_{A,max}$.

Intervention introduces biomass to the system and affects the transition terms in the matrix. $J_{temp_t}$ and $A_{temp_t}$ represent biomass added as temporary protected juveniles in the spring and removed as adults in the fall after they have released their spores. Introduced biomass of type "Jtemp" is completely protected from grazing and endures reduced mortality compared to juveniles on the benthos (McHugh et al., 2025). In our model, these temporary protected juveniles grow into adults with maturation rates $m_t$ and growth factors $\gamma_{A_t}$, same as their natural counterparts, but are not grazed ($g_{Jtemp} = 0$) and do not compete for light and space ($\beta_{JA_{temp}} = 0$). As adults, they are also protected from grazing ($g_{A_{temp}} = 0$) and mortality ($d_{A_{temp}} = 0$), while their breakage and fecundity are the same as their natural counterparts'. We do not include them in our count of the population's natural juvenile ($J_t$) and adult biomass $\sum_{a=1}^{a_{max}} A_t^a$. Restoration practitioners collect the adults, represented by $A_{temp_t}$, back from the system in the fall, after these temporary "outplants" have released their spores. We capture this in the model by zeroing out the $A_{temp_t}$ element in $\vec{X}_t$ at the end of every fall season. Introduced biomass of types "M", "J", and "Dssnl" experience the same demographic parameters as their natural counterparts.

#### *Landsat biomass estimates*

We downloaded Landsat kelp cover data from (kelpwatch.org) for the "Northern California" (San Francisco Bay to Oregon border) and "Southern California" (Santa Barabara to Mexico border) regions selected from the website's (Location) dropdown menu (Bell, Tom W et al., 2026). The satellite imagery cannot distinguish species, but due to their known ranges, we can reasonably assume that data from "Northern California" represent bull kelp and data from "Southern California" represent giant kelp.

We first converted the canopy area to total biomass in each region by multiplying canopy area from the Landsat data ($m^2$) with estimates of biomass density ($kg/m^2$) for the respective species (Rassweiler et al., 2025; Stekoll et al., 2006). To make the total biomass estimates comparable between the two regions, we divided each by their maximum canopy area from the Landsat time series data. Finally, we scaled the resulting $kg/m^2$ estimates up to 10,000$m^2$.

### *Environmental dynamics*

For each shared socio-economic pathway (SSP) climate projection from the GFDL-ESM4 model prepared for CMIP6, we classified every year from 2000-2100 as "good", "neutral", or "bad" according to the following rule: If at least 3 months of the year had projected sea surface temperature (SST) $>$ the 90th percentile of SST for the respective seasons over a 30-year rolling window, then we classified this year as "bad". If at least 3 months of a year had SST $<$ the 10th percentile of SST for the respective seasons over a 30-year rolling window, then we classified it as "good". We labelled the remaining years as "neutral". For all scenarios, we confirmed that no year satisfied the criteria for both "good" and "bad" years. We calculated frequencies of "good", "neutral", and "bad" years in the labelled SSP projections to estimate our year-type probability parameters $p_{good}$, $p_{neutral}$, and $p_{bad}$.

Our approach to classifying years as either "good", "neutral", or "bad" enables defining "good" and "bad" years together and consistently across scenarios, but misses the nuance that coastal waters can experience both stronger upwelling and warming. Brady et al. (2017) show that $\sim$50% of climate models within their ensemble predict at least 50% increase in springtime vertical velocity along the northern California coast under RCP-8.5 by 2100. Because we represent "good" years by cold temperature instead of upwelling velocity directly, we potentially underestimate the frequency of "good" nutrient-rich periods for kelp, even while SST warms over time.

The rolling climatology window approach accounts for the species of interest acclimating or adapting to changing climate and implicitly assumes a rate for this adaptation (Logan et al., 2014). A shorter window implies faster adaptation (i.e., the threshold at which kelp is physiologically stressed increases with the rate of ocean warming), while a longer window implies slower adaptation. We tested window lengths of 20, 30, and 40 years, and chose the intermediate window length which produced a reasonable range of outcomes across scenarios (i.e., for both species, frequency of simulations with persistence after 120 years was lower under more intense climate scenarios (Fig S10). The lack of robust data on the duration and magnitude of bull and kelp germ banks in the field complicates our certainty in the realized generation time and thermal adaptation rates for both species.

We calibrated year-type effects on kelp demographic parameters such that the majority of 100 simulations for both species showed kelp persistence at high biomass in the “historical” climate scenario, fewer simulations showed persistence in the SSP245 scenario, and most simulations resulted in population collapse in the extreme SSP585 climate scenario (Fig S10). We see an opportunity for future studies to take a more data-forward approach to characterizing the species’ comparative vulnerabilities to climate-related stress.

### Figure captions

**Figure S1: Density dependent recruitment**

Juvenile biomass plotted against microscopic biomass in the previous season. Data are from a single simulation with “historical” climate scenario, “low” grazing, and no intervention. Bull kelp (A-D) and giant kelp (E-F). A) and E) spring recruitment. B) and F) summer recruitment. C) and G) fall recruitment. D) and H) winter recruitment.

**Figure S2: Grazing on different kelp life stages**

Grazing rates on each life stage as a function of drift kelp abundance. Data are from a single simulation with "historical" climate scenario, "low" grazing, and no intervention.

**Figure S3: Burn-in and sensitivity to initial conditions**

Summer adult biomass over time. Each line represents one simulation from a unique initial biomass. All simulations are under "low" grazing, no intervention, and a deterministic "neutral" year climate scenario. A) Bull kelp. B) Giant kelp.

**Figure S4: Effects of life stages at introduction on outcomes, across grazing intensities**

Box plots showing distributions of outcome metrics across sets of introduced life stages (horizontal axes), climate scenario (color), and grazing intensity (columns). Top: bull kelp. Bottom: giant kelp. Each box plot, except for those of type "NONE", summarizes 90 data points. Each data point represents the mean of 100 stochastic simulations of a particular intervention case, i.e., combination of intervention cue, threshold, effort, type, wait time, and climate scenario. Dotted red lines indicate the mean of the respective outcome metric across 100 stochastic simulations in the "historical" climate scenario without intervention.

**Figure S5: Relative importance of decisions and biophysical attributes, using percent instead of percentile threshold**

Importance of variables for the following outcome metrics: A) population persistence, B) summer adult biomass, C) fall microscopic biomass. Importance is measured as the % decrease in MSE when the respective variable is removed from a random forest regression model predicting the outcome metric. Teal-green and black checkerboard labels grazing intensity, a variable that is both an intervention decision and a feature of the surrounding biophysical context.

**Figure S6: Simulated age structure**

Box plots showing biomass distributions in each age class of the adult (A and C) and microscopic (B and D) stages. Each box plot summarizes the final summer adult biomass or final fall microscopic biomass in an age class across 100 simulations. Yellow-green box plots show the results of 500$kg$ additions of temporary protected juveniles with 3 year wait-time, adult biomass cue, and percentile threshold of 0.1 for bull kelp and 0.05 for giant kelp. All simulations used the “low” grazing scenario. Bull kelp age structure (A and B). Giant kelp age structure (C and D).

**Figure S7: Effort scatterplot colored by percentile threshold**

Summer adult biomass (A-C, G-I) and population persistence (D-F, J-L) plotted against total intervention effort, across “low” (A, D, G, J), “medium” (B, E, H, K) and “high” (C, F, I, L) grazing scenarios. Bull kelp intervention outcomes (A-F). Giant kelp intervention outcomes (G-L). The maximum value on the horizontal axis indicates 120 years of 100% effort. Each point represents the mean of 100 stochastic simulations of a particular intervention case, i.e., combination of intervention cue, threshold, effort, type, wait time, and grazing and climate scenarios. Dotted red lines indicate the mean across 100 stochastic simulations of each outcome metric in the “historical” climate scenario without intervention. Points are colored by percentile threshold and shaped by life stages at introduction.

**Figure S8: Wait time and threshold correlations**

Summer adult biomass plotted against wait time (A-C, G-I); Persistence probability plotted against wait time (D-F, J-L); Summer adult biomass plotted against percent threshold (M-O, S-U); Persistence probability plotted against percent threshold (P-R,

V-X). Correlation coefficients are printed in blue on each panel. Data are from simulations across the full spectra of intervention decisions.

**Figure S9: Demographic parameter calibrations by year-type**

Frequency distributions of final summer adult biomass from 100 simulations in the “historical” (A and D), SSP245 (B and E) and SSP585 (C and F) climate scenarios. Bull kelp (D-F). Giant kelp ( A-C). Purple lines represent the 1st and green lines represent the 10th percentile of summer adult biomass in “historical” simulations.
