## Supplemental Figures for "Restoration decisions in highly variable systems: Whether, when, and how to enhance kelp"

S1

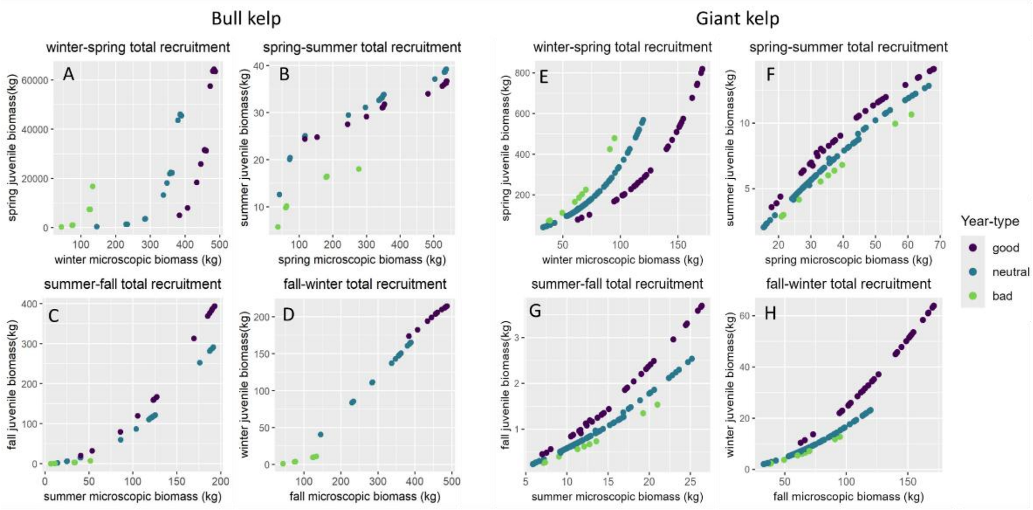


S2

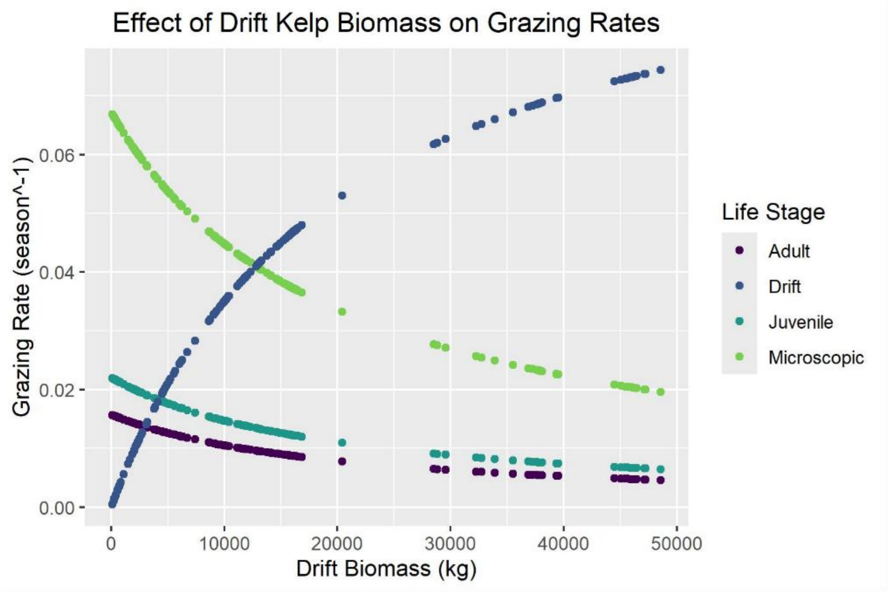

S3

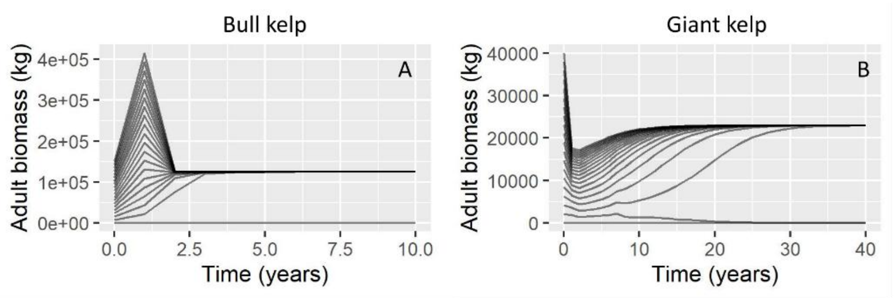
Bull kelp
Giant kelp
A
B
Adult biomass (kg)
4e+05
3e+05
2e+05
1e+05
0e+00
0.0
2.5
5.0
7.5
10.0
Time (years)
40000
30000
20000
10000
0
0
10
20
30
40
Time (years)

S4

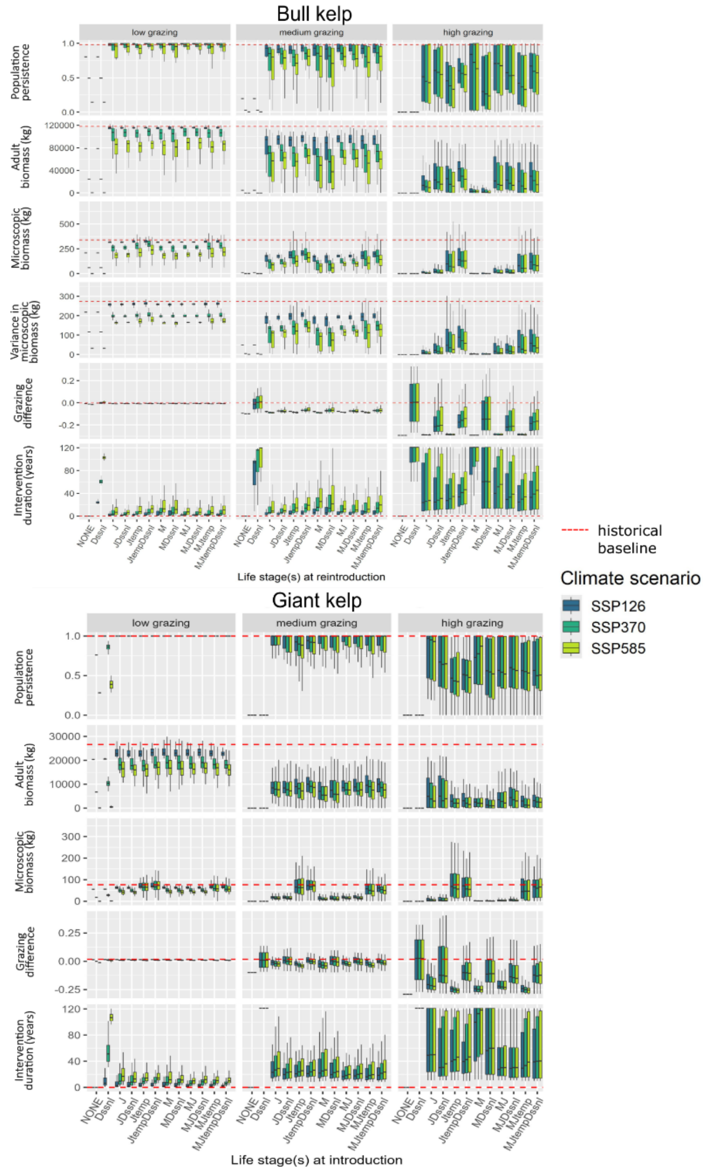
Bull kelp
low grazing
medium grazing
high grazing
Population persistence
Adult biomass (kg)
Microscopic biomass (kg)
Variance in microscopic biomass (kg)
Grazing difference
Intervention duration (years)
NONE
Dssni
J
JDssni
Jtemp
JtempDssni
M
MDssni
MJ
MJDssni
MJtemp
MJtempDssni
Life stage(s) at reintroduction
historical baseline
Climate scenario
SSP126
SSP370
SSP585
Giant kelp
Life stage(s) at introduction

S5

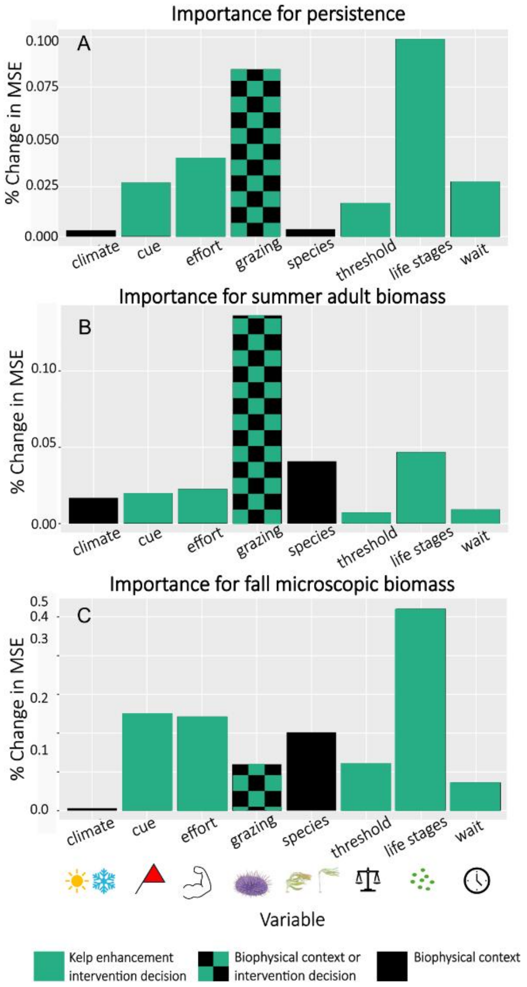
Importance for persistence
A
% Change in MSE
0.100
0.075
0.050
0.025
0.000
climate
cue
effort
grazing
species
threshold
life stages
wait
Importance for summer adult biomass
B
% Change in MSE
0.10
0.05
0.00
climate
cue
effort
grazing
species
threshold
life stages
wait
Importance for fall microscopic biomass
C
% Change in MSE
0.5
0.4
0.3
0.2
0.1
0.0
climate
cue
effort
grazing
species
threshold
life stages
wait
Variable
Kelp enhancement intervention decision
Biophysical context or intervention decision
Biophysical context

S6

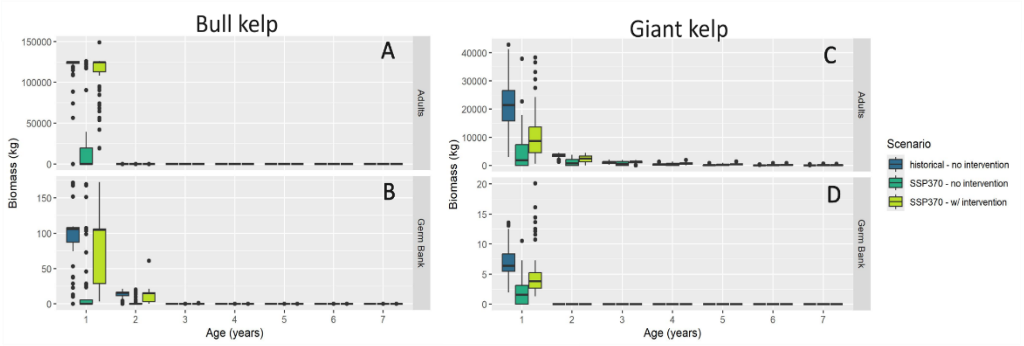
Bull kelp
A
B
Biomass (kg)
150000
100000
50000
0
150
100
50
0
Adults
Germ Bank
Age (years)
Giant kelp
C
D
Biomass (kg)
40000
30000
20000
10000
0
20
15
10
5
0
Age (years)
Scenario
historical - no intervention
SSP370 - no intervention
SSP370 - w/ intervention

S7

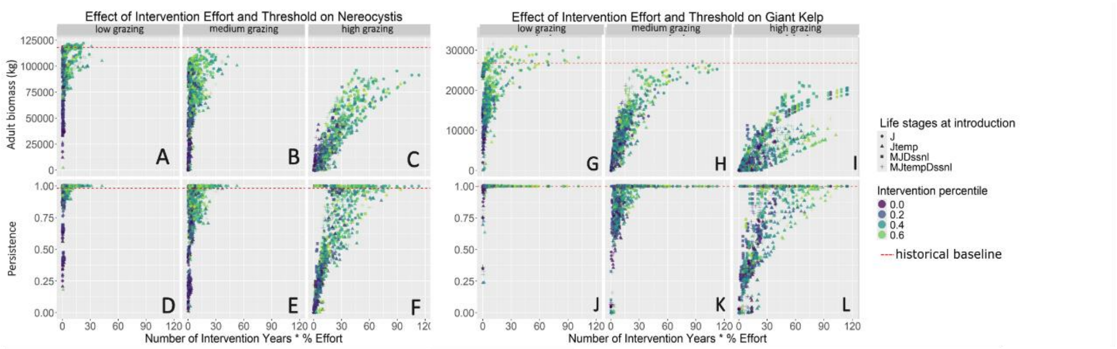
Effect of Intervention Effort and Threshold on Nereocystis
Effect of Intervention Effort and Threshold on Giant Kelp
low grazing
medium grazing
high grazing
Adult biomass (kg)
Persistence
Number of Intervention Years * % Effort
Life stages at introduction
J
Jtemp
MJDssnl
MJtempDssnl
Intervention percentile
historical baseline

S8

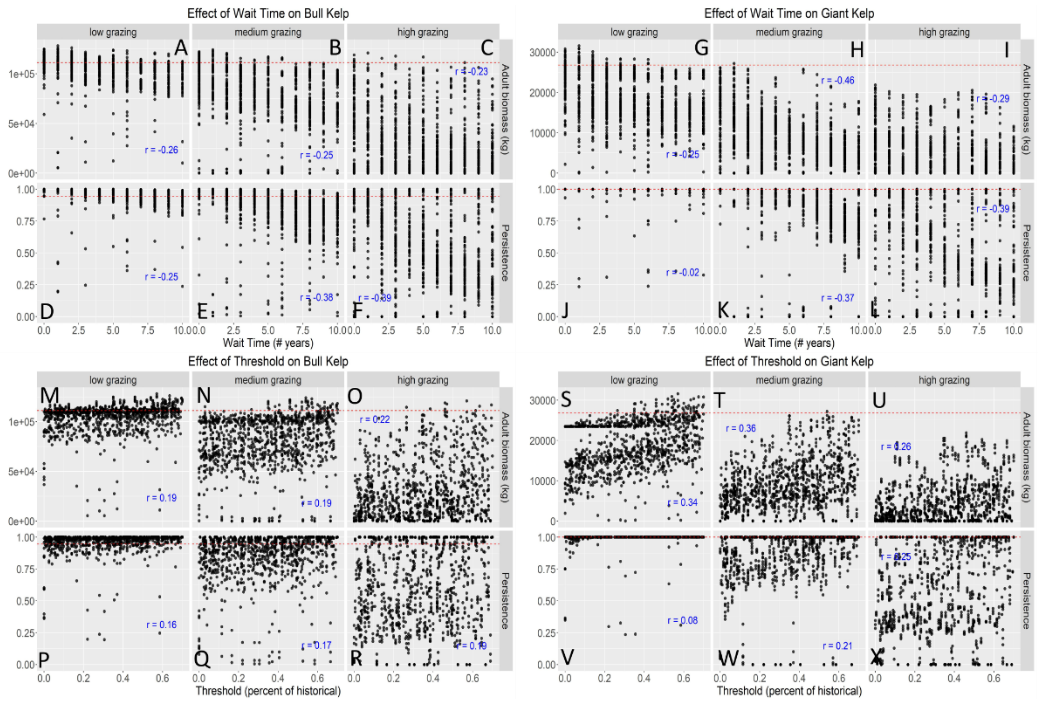
Effect of Wait Time on Bull Kelp
Effect of Wait Time on Giant Kelp
Effect of Threshold on Bull Kelp
Effect of Threshold on Giant Kelp
low grazing
medium grazing
high grazing
Adult biomass (kg)
Persistence
Wait Time (# years)
Threshold (percent of historical)

S9

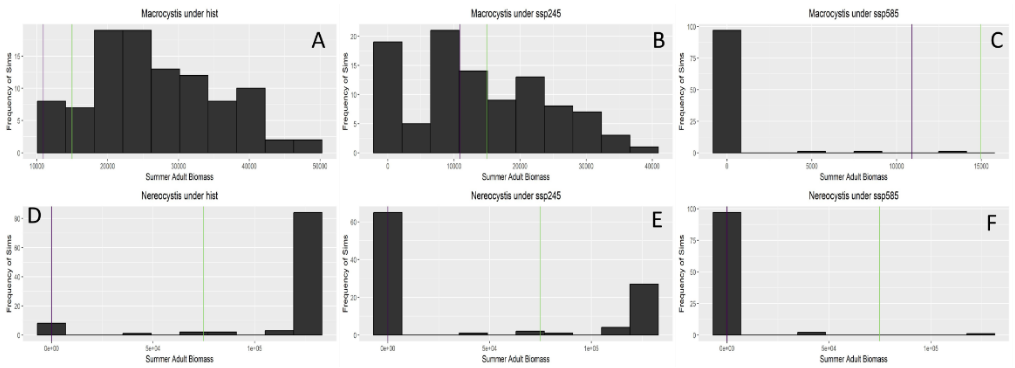
Macrocystis under hist
Macrocystis under ssp245
Macrocystis under ssp585
Nereocystis under hist
Nereocystis under ssp245
Nereocystis under ssp585
Frequency of Sims
Summer Adult Biomass
